# scATrans: annotating single-cell differential expression as transcription- or stabilization-weighted using unspliced RNA

**DOI:** 10.64898/2026.08.03.740741

**Authors:** Zhao Li, Aaron W. James, Shengxuan Li

**Affiliations:** Department of Pathology, Johns Hopkins University, Baltimore, MD, 21205, USA

**Keywords:** single-cell RNA-seq, differential expression, unspliced RNA, nascent RNA, transcription versus mRNA stabilization, mechanism annotation, metabolic labeling, pseudobulk, gene-set analysis

## Abstract

Single-cell differential expression (DE) reports changes in mature mRNA abundance, but the same fold-change can reflect faster synthesis or slower decay. Metabolic labeling resolves this ambiguity but is costly and cannot be applied retrospectively to the vast majority of published scRNA-seq. scATrans closes this gap using layers every standard pipeline already generates: from DE-selected genes, a reference-corrected unspliced residual annotates each change as transcription- or stabilization-weighted, with no additional experiment. Benchmarked against metabolic-labeling systems with independent kinetic ground truth, the residual separates the two mechanisms at matched mature abundance (ROC-AUC 0.68–0.74, full-length NASC-seq2 K562; 0.59–0.63, 3′ scEU-seq RPE1); effect size scales with intron capture, not model complexity, and explicit kinetic fitting adds nothing over the static contrast. Per-gene, the residual recovers the classical bulk exon–intron contrast (EISA); what scATrans adds is the inference layer single-cell reanalysis actually needs—DE-defined membership, gene-structure correction, a capture-regime reliability pre-flight, induction-matched testing, and a permutation-calibrated program score—so that confident calls are reserved for where the data support them: gene programs, not single genes. Applied to standard 10x data with no labeling, scATrans recovers textbook post-transcriptional biology: a curated AU-rich-element program is called stabilization-weighted in LPS-stimulated PBMCs (confirmed by per-donor pseudobulk DE in an independent four-donor cohort), while a glucocorticoid-response program is called transcription-weighted in dexamethasone-treated A549 cells— opposite mechanisms recovered from unlabeled counts. scATrans turns any spliced/unspliced-resolved DE table into a mechanism-typed one, retrospectively and at scale.

**Availability and implementation:** scATrans requires Python ≥3.9, interoperates with the scverse ecosystem (AnnData, Scanpy), and is released under the Apache-2.0 license. Install with pip install scatrans or from Bioconda. Documentation and tutorials: https://scatrans.readthedocs.io. Analyses in this manuscript use software version 0.10.9.

## Introduction

Single-cell differential expression (DE) reports changes in mature mRNA abundance ^1^, but the same fold-change can arise from faster transcription or slower decay— mechanisms that imply different regulatory nodes and different intervention points. In LPS-stimulated immune cells, for instance, NF-κB drives transcription while ARE-mediated stabilization prolongs cytokine mRNA ^2,3^; DE detects both yet distinguishes neither.

Metabolic labeling resolves the ambiguity: a nucleoside-analog pulse marks nascent RNA, enabling direct measurement of synthesis and decay at single-cell resolution ^4–7^. Yet labeling requires timed pulses, specialized chemistry, and dedicated libraries—costs that preclude retrospective application to the millions of unlabeled scRNA-seq samples already in public repositories ^8^.

An alternative already exists in every standard pipeline: spliced and unspliced counts, produced at no extra cost ^9,10^, where unspliced reads enrich for nascent transcripts. RNA-velocity methods exploit this layer to extrapolate trajectories, but identifiability and interpretability remain contested ^11,12^. A simpler, static use—annotating DE results as transcription- or stabilization-weighted without fitting kinetic models—has not been systematically developed.

The underlying statistic is not new. Bulk exon–intron split analysis (EISA) already separates transcriptional from post-transcriptional change by contrasting intronic and exonic fold-changes ^13^, and—as we confirm below—the per-gene signal in single-cell data is equivalent. What is missing is the inference layer: DE-defined gene membership, gene-structure correction, a capture-regime reliability check, and permutation-calibrated program-level testing. scATrans supplies that layer, positioning the exon–intron contrast for routine single-cell mechanism annotation.

scATrans implements this framework as an open-source Python package. From DE-selected genes it computes a gene-structure-corrected unspliced residual; at matched mature abundance, excess unspliced signal indicates transcription-weighted change, deficit indicates stabilization. Per-gene resolution is limited, so confident calls are made at the program level via permutation-calibrated testing. Below we validate the residual against metabolic-labeling ground truth, confirm equivalence with the bulk exon–intron contrast, and demonstrate that the framework recovers known post-transcriptional biology—ARE-mediated stabilization in LPS-stimulated PBMCs, transcriptional induction in glucocorticoid-treated cells—from standard 10x data without any labeling.

## Results

### The residual separates transcription-from stabilization-weighted abundance across metabolic-labeling systems

We benchmarked the residual against three metabolic-labeling datasets with independent kinetic ground truth: scEU-seq RPE1 pulse–chase (3′), scEU-seq mouse intestinal organoid (3′), and full-length NASC-seq2 K562 ^7^—each providing per-gene degradation rates (γ) from labeled/unlabeled kinetics, independent of the spliced/unspliced layers. At matched mature abundance, the residual separated fast-degrading (stabilization-weighted) from slow-degrading (transcription-weighted) genes with ROC-AUC 0.59–0.63 on 3′ RPE1 and 0.68–0.74 on full-length K562 (Fig. 1A). The sparser 3′ organoid yielded AUC ≈0.54, near chance—intron capture, not model complexity, limits resolution. Gene-structure residualization removed the length and GC bias otherwise present in the raw score (Fig. 1B; Fig. S3)

The residual captures transcription-rate information independent of abundance. Controlling for log abundance, partial correlation between residual and log γ was +0.148 (Fig. 1D); the DE analogue is zero by construction. We defined transcription-versus stabilization-weighted genes as top versus bottom γ tercile within each abundance decile (n = 2,549; abundance-matched Mann–Whitney p = 0.46). The residual reached ROC-AUC 0.627 while DE showed no discriminative power (0.494); within-decile pooling gave 0.624 (Fig. 1A). For reference, the oracle upper bound—using true labeled unspliced counts—is ≈0.68. Gene-structure residualization costs some signal but does not eliminate it: 0.615 under package defaults, 0.591 after full length/intron correction (Fig. 1B). Length control also matters for interpretation: it reduces genome-wide ARE-density and chromatin associations to length artifacts (Fig. S3). After length correction, partial correlation with γ remained positive (ρ = +0.119) while correlation with gene length collapsed from +0.277 to +0.083.

Mature abundance is anti-aligned with turnover: Spearman ρ(spliced, γ) = −0.457 (AUC 0.213 for recovering high-turnover genes). At steady state A = k/γ, so high-abundance genes by DE are disproportionately stable. Unspliced counts point the other way, though weakly (ρ = +0.037; Fig. 1C). The residual strengthens this ranking: Spearman ρ with γ = +0.14 (Table 1), partial ρ = +0.148 controlling for abundance (Fig. 1D), matched-mechanism AUC 0.627 (Fig. 1A). Abundance systematically mis-orders turnover; the residual corrects it.

Cross-system comparison confirmed that effect size tracks intron capture. The scEU-seq intestinal organoid recovered the same direction only weakly (matched AUC ≈0.54; Table 1)—a low-signal system serving as negative control for heavily parameterized estimators rather than independent validation. The NASC-seq2 K562 dataset, using orthogonal labeling chemistry (4sU conversion) with full-length reads and substantially better intron capture, gave the strongest result: matched-abundance AUC 0.742 (0.716 under package defaults, 0.681 after length/intron correction; Fig. 1B) versus DE 0.496, with partial ρ(residual, γ | abundance) = +0.291 (n = 3,040). Leave-one-decile-out and bootstrap audits confirmed a broad-based result (95% CI 0.730–0.755) rather than outlier-driven. The ordering—full-length K562 (0.74) > 3′ RPE1 (0.63) > sparse organoid (≈0.54)—scales with intron-capture quality across platforms and labeling chemistries. Where capture is adequate, the residual resolves a distinction that DE conflates.

**Figure 1.**
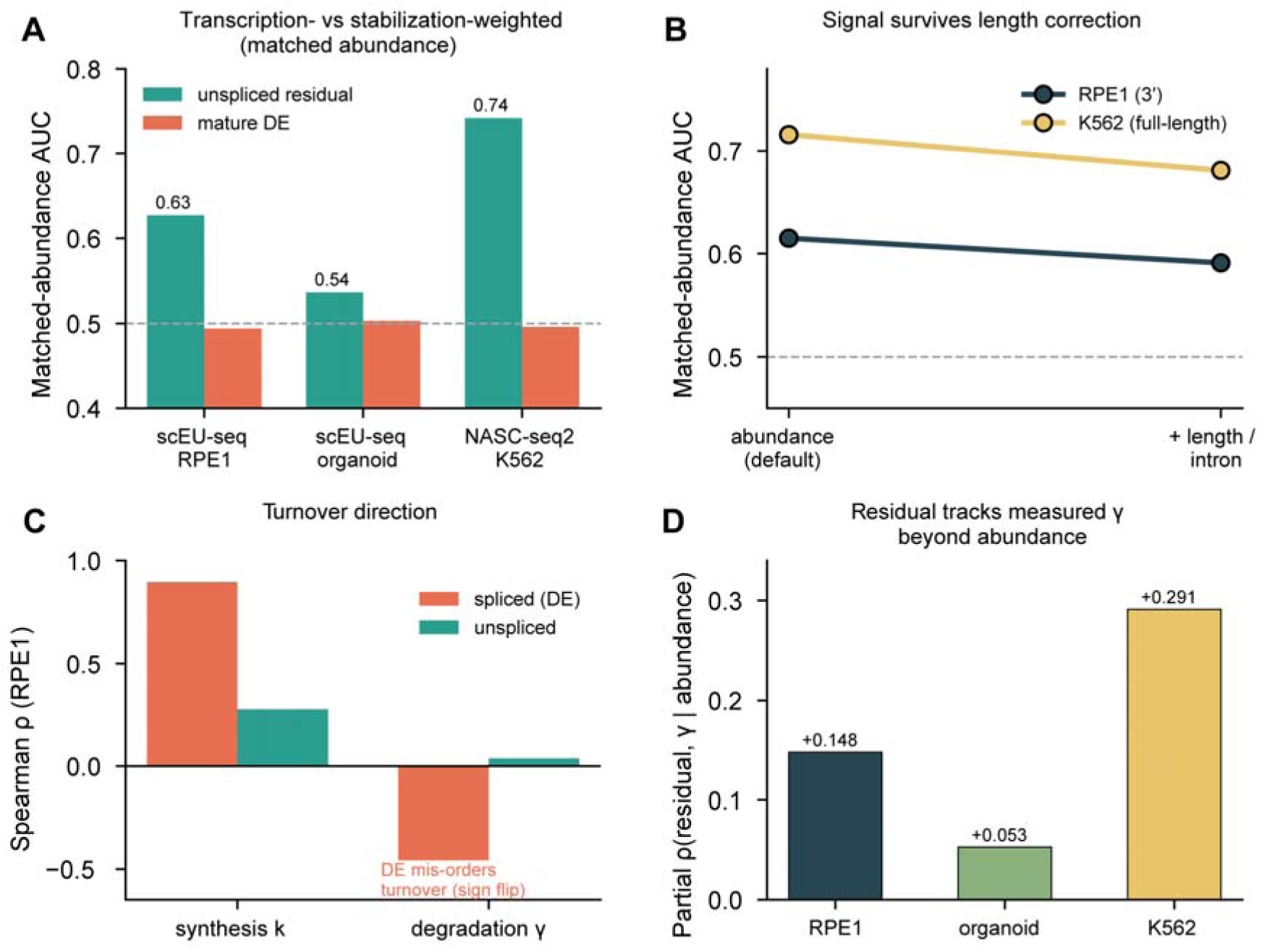
Specificity of the residual for transcription-versus stabilization-weighted expression. (**A**) Matched-abundance ROC-AUC of the residual versus mature DE across scEU-seq RPE1, scEU-seq organoid, and NASC-seq2 K562; DE is near chance in all three. (**B**) Matched-abundance AUC under package-default abundance residualization (K562 0.716, RPE1 0.615) and after full length/intron correction (0.681, 0.591); headline values without structure residualization are in (A) (0.742, 0.627). Residualization reduces but does not eliminate the residual–DE gap. (**C**) On RPE1, spliced abundance is anti-correlated with measured turnover (sign flip); unspliced counts track high-turnover genes in the correct direction, weakly (ρ = +0.037). (**D**) Partial correlation between the residual and degradation rate γ, controlling for abundance (RPE1 +0.148, organoid +0.053, K562 +0.291); the DE analogue is zero by construction. Failed orthogonal validators in Fig. S3.

### The static residual attains kinetic-model performance without kinetic assumptions

We compared five predictors against measured-γ ground truth (Table 1): scATrans residual, raw u/s ratio, EISA intron-level contrast, steady-state velocity γ, and scVelo dynamical-model γ. scEU-seq RPE1 and organoid supported all five (per-cell layers + independent γ); NASC-seq2 K562 contributed only the first three, as velocity estimators require cross-cell regression.

The raw u/s ratio correlated more strongly with γ than the residual (Spearman +0.42 vs +0.14, RPE1)—expected, since residualization trades raw correlation for abundance-independence. At matched abundance the residual led (AUC 0.627 vs 0.613).

Against the exon–intron baseline ^13^, the residual and EISA are equivalent as per-gene scores: matched AUC 0.742 for both on K562; 0.627 vs 0.618 on RPE1. The bulk intronic contrast already captures most per-gene information in these layers—we claim no improvement on the statistic itself. What scATrans adds is the inference framework: DE-coupled membership, gene-structure residualization (without which ranking partly reflects length; Fig. 1B; Fig. S3), capture-regime pre-flight, induction-matched program testing, and a permutation-calibrated score that provides an empirically grounded zero. Kinetic fitting added nothing. Both velocity estimators performed at or below the static residual; the full dynamical model was weakest, falling below chance on the sparse organoid (Table 1). scVelo returns usable γ for only a few hundred genes; the residual covers thousands ^11,12^. We therefore treat the residual as a static contrast, not inferred rates.

That the framework, not the statistic, does the work is a testable claim—two negative results confirm it. Two per-gene scores that match or tie the residual on the γ benchmark fail at the program level: the single-sample intronic predictor and the transcription-change leg of EISA alone both miss the ARE stabilization call in the LPS case study below (Table S3). Per-gene equivalence does not guarantee program-level inference; the intervening steps—membership, residualization, induction matching, calibration—are where transfer is won or lost. The full exon–intron contrast does reproduce the call but cannot place it against an empirical zero; we return to this comparison below.

**Table 1.** Baseline comparison against measured degradation rate γ (scEU-seq RPE1, NASC-seq2 K562 and scEU-seq intestinal organoid). All predictors use matched-abundance AUC (chance = 0.50; bootstrap 95% CI; Methods); package-default and fully residualized values in Fig. 1B. Spearman is within-system only (RPE1/organoid γ are absolute pulse–chase rates; K562 γ is a relative labeling rate). Raw ratio can lead on Spearman yet trail on matched AUC because abundance drives much of that correlation. Velocity γ is undefined on K562 (single pseudobulk; both estimators require cross-cell regression). The organoid is a weak system (static predictors ≈0.53); dynamical γ falls below chance and is reported as caution, not support.

| System | Predictor (spliced/unspliced) | Spearman($\gamma$) | Matched-AUC (95% CI) | n |
| --- | --- | --- | --- | --- |
| RPE1 | scATrans static residual (matched-abundance) | **+0.14** | **0.627 (0.611–0.643)** | **5,098** |
| RPE1 | Raw unspliced-to-spliced ratio | +0.42 | 0.613 (0.602–0.626) | 5,098 |
| RPE1 | EISA intron level (exon–intron split; matched-abundance) | +0.04 | 0.618 (0.603–0.633) | 5,098 |
| RPE1 | RNA-velocity steady-state $\gamma$ (linear fit) | +0.39 | 0.596 (0.583–0.610) | 5,098 |
| RPE1 | RNA-velocity dynamical-model $\gamma$ (recover_dynamics)* | +0.21 | 0.584 (0.561–0.609) | 2,000 |
| K562 | scATrans static residual (matched-abundance) | **+0.32** | **0.742 (0.730–0.755)** | **6,078** |
| K562 | Raw unspliced-to-spliced ratio | +0.41 | 0.683 (0.673–0.694) | 6,078 |
| K562 | EISA intron level (exon–intron split; matched-abundance) | +0.31 | 0.742 (0.730–0.754) | 6,078 |
| organoid | scATrans static residual (matched-abundance) | +0.03 | 0.537 (0.521–0.554) | 4,694 |
| organoid | Raw unspliced-to-spliced ratio | +0.06 | 0.535 (0.522–0.549) | 4,694 |
| organoid | EISA intron level (exon–intron split; matched-abundance) | +0.04 | 0.537 (0.521–0.554) | 4,694 |
| organoid | RNA-velocity steady-state $\gamma$ (linear fit) | +0.07 | 0.536 (0.522–0.551) | 4,694 |
| organoid | RNA-velocity dynamical-model $\gamma$ (recover_dynamics)* | –0.14 | 0.427 (0.405–0.456) | 2,000 |

*Dynamical $\gamma$ reported for most favorable configuration (recover\_dynamics forced onto 3,000 highest-expressed ground-truth genes; 2,000 in extreme abundance-matched $\gamma$ terciles enter the comparison). scVelo default gene selection and canonical HVG pipelines fit fewer genes (160 and 517) and give chance-level $\gamma$ recovery (AUC 0.52, 0.50); 0.584 is the dynamical model's best case.*

### The residual peaks early after stimulation and decays as mature mRNA accumulates

If the residual tracks nascent RNA, its signal should peak before transcriptional changes propagate into the mature pool, then decay. Two stimulus time courses confirmed this prediction. On scNT-seq neurons, residual AUC fell monotonically from 0.70 (15 min) to 0.48 (120 min) as DE strengthened (Fig. 2A). On sci-fate A549 dexamethasone, the residual decayed from 0.65 (2 h) toward chance (≈0.56 by 10 h) while DE rose (Fig. 2B). The residual does not outperform DE for gene discovery—its value lies in mechanism direction. Acute responses are therefore best read early or at program level.

**Figure 2.**
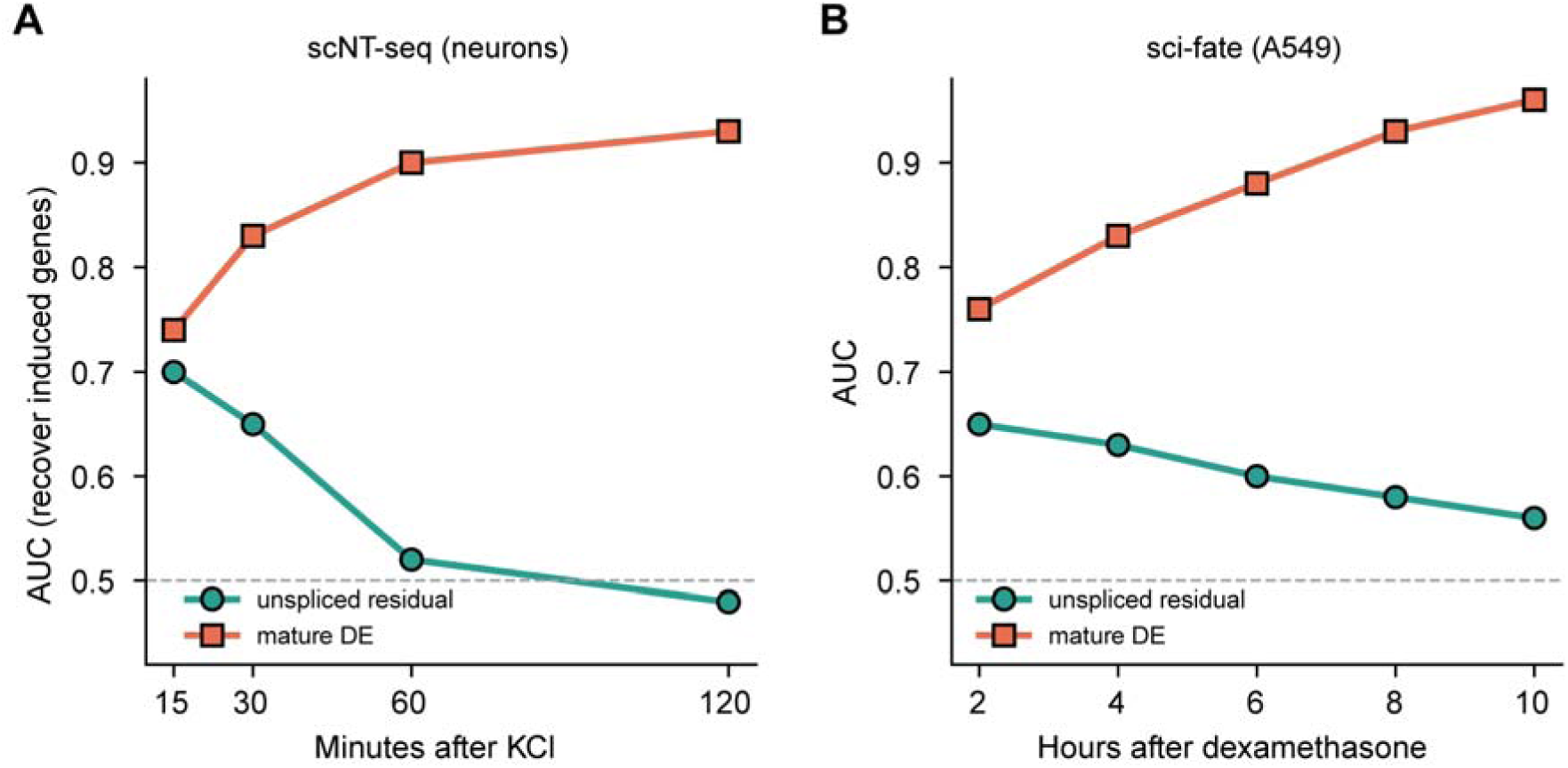
Time-dependent residual performance on metabolic-labeling stimulus responses. (**A**) scNT-seq KCl neurons: residual AUC decays toward chance as mature DE strengthens. (**B**) sci-fate A549 dexamethasone: same pattern on an independent platform. The residual is informative early but does not exceed DE for recovering induced genes.

### Program-level mechanism calls recover ARE-linked stabilization biology in real 10x scRNA-seq

We tested whether the framework recovers known biology on unlabeled droplet data. Public 10x v3.1 PBMCs (resting vs 4 h LPS; 15,449 cells; GSE226488) were reprocessed with STARsolo Gene+Velocyto (GRCh38). Global unspliced fraction (0.327) passed the reliability pre-flight (Fig. 3A). Design caveats apply: one library per condition, single donor in the stimulated arm, cell-level DE without biological replication—p-values are exploratory ^14–16^. This is a biological-plausibility test: does textbook ARE biology emerge? Polarity is later confirmed in an independent four-donor series.

LPS induces inflammatory cytokines via two textbook mechanisms: NF-κB-driven transcription and ARE-mediated mRNA stabilization (notably TTP/ZFP36 targets) ^2^. These form a natural positive contrast. We tested whether a curated ARE/ZFP36 set reads as more stabilization-weighted than a primary NF-κB transcriptional set among LPS-responsive genes (Table S2; curation in Methods S5). Lists are disjoint by construction; dual-regulated genes were assigned to at most one. The ARE set comprised 21 curated symbols → 19 in the index → 17 with finite transcription_support → 16 in the DE-selected program; ZFP36 itself was not induced (log2FC = −0.67) and is excluded.

A key confounder: strongly induced genes accumulate mature mRNA by 4 h, so induction strength can lower unspliced excess independently of mechanism. Across the induced universe this coupling is weak (ρ = +0.19, n = 1,943), so induction alone does not explain program position—but induction-matched tests (comparing residuals among genes at similar log2FC) provide a direct exclusionary check. The matched test, not the raw program mean, is the primary analysis.

Within the DE-selected program (padj < 0.05, log2FC ≥ 1), the ARE set was called stabilization-weighted (n = 16; mean transcription_support −17.45, median −8.46, vs background +0.54/+0.29; Mann–Whitney p = 1.6×10 ; Fig. 3B). Both statistics are reported because the distribution is heavy-tailed: CCL4 and CCL3 pull the mean, but the median confirms the shift holds for typical ARE genes.

The NF-κB set sat above ARE but was not a positive absolute call (mean −0.38, median −0.76, n = 10)—at 4 h it lay slightly below background. This reflects the induction confound: strongly induced genes accumulate mature mRNA, displacing uncalibrated means downward. Absolute placement therefore requires permutation calibration: a 1,000-run condition-label shuffle supplied the gene set’s null expectation (≈ −2.9); calibrated displacement is reported in Fig. 3F. Conclusion: a curated ARE program reads as stabilization-weighted relative to background on standard 10x data.

The call survived induction matching (Fig. 3C). OLS on the induced universe (n = 1,943; 17 ARE members) regressing winsorized transcription_support on log2FC + ARE indicator gave β_ARE = −4.97 (SE 0.60, p = 1.6×10 ¹); log2FC was non-significant (β = +0.04, p = 0.21). On this universe’s scale (winsorized SD = 2.45), β_ARE ≈ −2 SD, moving a typical gene from median to ∼2nd percentile. Without winsorization, β = −16.8—the reported value is conservative. Nearest-neighbor matching (each ARE gene paired with five non-ARE genes at similar log2FC) confirmed: median support deficit −6.38 (Wilcoxon p ≪ 0.001). The stabilization call is not an artifact of stronger ARE induction.

Four independent checks agreed. Pre-ranked GSEA placed the ARE/ZFP36 set at the stabilization pole (NES = −2.76, FDR < 0.001) but not the NF-κB set (NES = −1.30, FDR = 0.15; Fig. 3D). Leave-one-out refitting kept β_ARE within [−5.32, −4.75], all p < 2×10□¹□, including removal of the extreme outlier CCL4; alternate curations gave −4.89 to −5.74 (Supplementary Methods S5). Within 3,893 gated monocytes—the lineage expressing these cytokines—the signal strengthened (β_ARE = −8.13, p = 7.7×10□³²; NF-κB weaker at β = −1.89; Table S4), ruling out cell-composition effects. A 1,000-run condition-label permutation confirmed the deficit against the gene set’s own shuffled expectation (p = 0.0010).

Permutation calibration supplies zero-referenced placement. Each program mean is compared with its expectation over 1,000 condition-label shuffles on the identical DE table and program membership, so all numbers below refer to one universe (803 DE genes; ARE n = 16, NF-κB n = 10). The ARE null expectation was −2.94 (SD 0.13) rather than zero; its observed mean of −17.45 gave a calibrated displacement of −14.52 (permutation p = 0.0010; no shuffle reached the observed deficit). The NF-κB set moved as calibration predicts: before calibration it sat at −0.38, its null expectation was −0.15, and after calibration it sat at −0.23—a quarter of one genome-wide unit from zero—while ARE stayed at the stabilization pole (Fig. 3F). The uncalibrated NF-κB displacement was small to begin with and rests on ten genes, so a transcription-weighted program with a large uncalibrated offset would test the procedure harder. What the pair establishes is that calibration does not remove signal: applied to two programs on the same universe, it returned one to the neighborhood of zero and left the other’s displacement intact. Displacements are not converted to z-scores because the null SD measures stability of the structural offset across shuffles, not sampling spread of the program mean.

The harder test comes from the glucocorticoid case. The identical procedure applied to the GR program (sci-fate A549, dexamethasone 2 h; n = 20) gave an observed mean of +0.522, null expectation +0.430 (SD 0.014), and calibrated displacement +0.0 92 (permutation p = 0.0010; Fig. S1D). Here 82% of the raw positive offset was structural, yet what remained stayed on the transcription side—the mirror of the ARE result on a different platform, cell type, and stimulus. The two null expectations have opposite signs (−2.94 for ARE, +0.43 for GR) despite both programs being short and low-intron: the structural offset is not a fixed function of gene architecture and must be estimated per gene set and dataset.

EISA reproduced the ARE stabilization call at the same standardized effect as the residual (−1.85 versus −1.87 winsorized SD on the common gene universe; Table S3). The two scores are weakly correlated and confounded by induction in opposite directions, so agreement is corroboration, not redundancy. What they do not share is a zero: EISA can rank ARE below NF-κB but cannot place NF-κB against an empirical null. Ordering comes from both scores; zero-referenced placement only from the calibrated residual. Two negative per-gene predictors that lose the call are in Table S3.

The axis extends to transcription-weighted programs. In sci-fate A549 cells treated with dexamethasone (GSE131351), a curated GR program (FKBP5, TSC22D3/GILZ, PER1, DUSP1, SGK1, DDIT4, and related genes; n = 20) read as transcription-weighted. At 2 h, GR targets showed higher transcription_support than background (Mann–Whitney p = 0.011 one-sided, p = 0.022 two-sided) with positive induction-matched coefficient (β_GR = +0.92, p = 0.010; nearest-neighbor median +0.84, 65% positive), decaying to β ≈ 0.6– 0.7 by 4–8 h in line with the residual’s nascent time decay (Fig. 2, Fig. S1). This is the opposite polarity to the ARE program and demonstrates both directions of the axis on real perturbations. The GR effect is smaller than β_ARE; the two are not directly comparable since they are estimated in different gene universes with different support distributions, and stabilization calls separate more cleanly than transcription calls in a heavy-tailed distribution. Because sci-fate measures new RNA, we confirmed these GR genes were transcriptionally induced (new-RNA log2FC ≈ +1.5–1.8); the residual recovered the program without that label.

**Figure 3.**
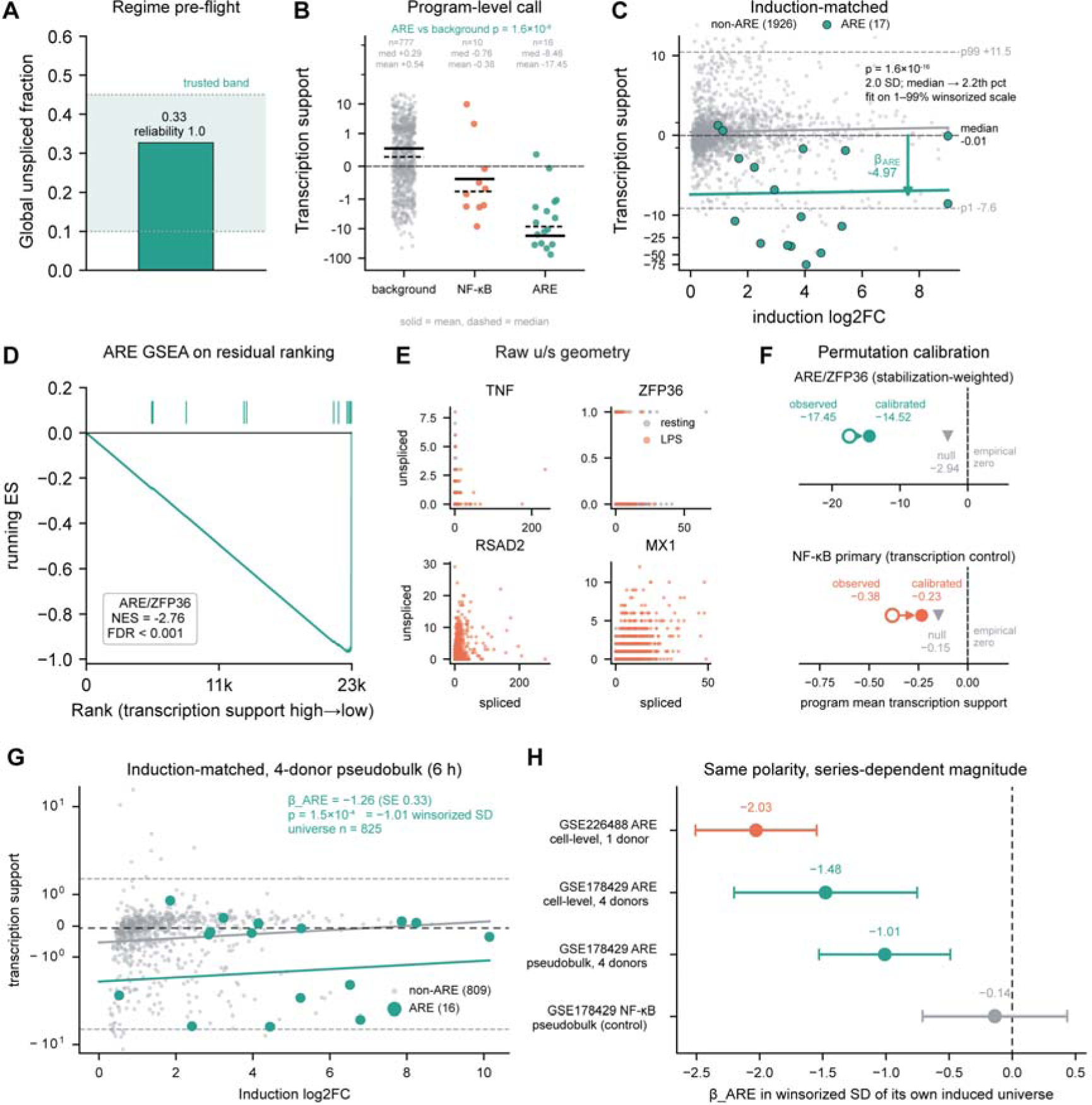
LPS–PBMC mechanism partition, ARE biology, and four-donor replication. (**A**) Regime pre-flight: global unspliced fraction 0.33 (reliability 1.0) on 15,449 cells, inside trusted band. (**B**) Program-level call (background n = 777, NF-κB n = 10, ARE/ZFP36 n = 16; solid = mean, dashed = median; symlog axis). ARE mean −17.45 / median −8.46 versus background +0.54 / +0.29 (Mann–Whitney p = 1.6×10□□); NF-κB −0.38 / −0.76, above ARE but slightly below background at 4 h (per-gene induction confound in Fig. S4A). Leave-one-out refits confirm the call does not rest on CCL4 or CCL3 (Methods S5). (**C**) Induction-matched regression (n = 1,943; 17 ARE members): OLS of winsorized transcription_support on log2FC + ARE membership gives β_ARE = −4.97 (SE = 0.60, p = 1.6×10□¹□), 2.0 winsorized SD (median → ∼2nd percentile). (**D**) Pre-ranked GSEA: NES = −2.76, FDR < 0.001; ticks mark ARE ranks. (**E**) Raw unspliced-versus-spliced geometry for four exemplars (TNF, ZFP36, RSAD2, MX1), descriptive only. (**F**) Permutation calibration (same universe and membership as B): observed mean (open circle), null expectation over 1,000 shuffles (gray triangle), calibrated displacement (filled circle). ARE: −17.45 → −14.52, stays at stabilization pole; NF-κB: −0.38 → −0.23, near empirical zero. (**G**) Four-donor replication (GSE178429, 6 h): per-donor pseudobulk DESeq2 (n = 825; ARE n = 16; β_ARE = −1.26, SE = 0.33, p = 1.5×10 ; −1.01 winsorized SD). (**H**) Cross-series comparison on one scale: GSE226488 cell-level (−2.03), GSE178429 cell-level (−1.48), GSE178429 pseudobulk (−1.01), NF-κB control (−0.14, CI spans zero). Polarity replicates; magnitude is series-dependent. Induction evidence and donor-pooling rationale in Fig. S5.

To rule out design artifacts, we first tested whether the DE-dependent steps are sensitive to single-library, cell-level testing (pseudoreplication ^14,15^). We repeated the workflow on IFN-β–stimulated PBMCs from eight demuxlet-resolved donors (Kang et al., GSE96583 ^17^) under a replicate-valid design—per-donor pseudobulk DESeq2, eight biological replicates per condition (Fig. S2). Of 312 cell-level-selected induced genes, 310 (99.4%) were re-selected under pseudobulk DESeq2 ^18,19^; both lists enriched the same interferon/antiviral program, and the 558 additional genes from the replicate design carried weaker secondary immune signal rather than noise. Disagreement concentrated in significance ranking, where pseudoreplication acts: concordance AUC 0.97 by effect size but only 0.77 by signed significance. Critically, transcription_support is computed from spliced/unspliced layers alone and was numerically identical between designs (max |Δ| = 0 across 62,754 genes), so mechanism calls depend on gene membership—99.4% stable—not on DE p-values. Because IFN-β does not induce the ARE program, this replicates DE-invariance of the mechanism axis and stability of program membership, not the ARE effect size.

Because the Kang dataset uses an IFN-β stimulus lacking ARE induction, we subsequently validated the biological effect size using the Kartha LPS series. The ARE effect size replicated in this independent multi-donor LPS series. The Kang series cannot supply this test: its ARE members were not induced (median log2FC = −0.24; 3 of 21 curated symbols passing |log2FC| ≥ 1, versus 12 of 12 interferon-stimulated controls at median +4.66)—the wrong stimulus, not insufficient power. We therefore reprocessed the Kartha et al. LPS series (GSE178429) from raw FASTQ with the same STARsolo configuration, obtaining 13,438 cells, and analyzed under a replicate-valid design: per-donor pseudobulk DESeq2 with four biological replicates per condition.

Here the ARE program was genuinely induced—median pseudobulk log2FC rising from +1.16 at 1 h to +3.97 at 6 h, with 17 of 19 curated members passing |log2FC| ≥ 1 at 6 h (Fig. S5A)—and the induction-matched stabilization call reproduced at 6 h: β_ARE = −1.26 (SE = 0.33; p = 1.5×10) over 825 induced genes with 16 ARE members (Fig. 3G). Nearest-neighbor matching and the competitive test (Mann–Whitney p = 0.038) agreed in sign; the NF-κB set was undisplaced in the same fit (Fig. 3G, H; Table 2). The contrast is program-specific. Polarity is the primary claim; standardized magnitude is secondary.

Fig. 3H places both series on one winsorized-SD scale (cell-level single-donor −2.03 → cell-level multi-donor −1.48 → four-donor pseudobulk −1.01; NF-κB −0.14): polarity agrees while magnitude is roughly half that of the primary series. Series, time, and capture differences each act in the observed direction (Fig. 2; Fig. 1B). We treat polarity as replicated and magnitude as series-dependent. The 1 h point is directionally consistent only (17 genes under per-donor pseudobulk DE).

**Table 2.**
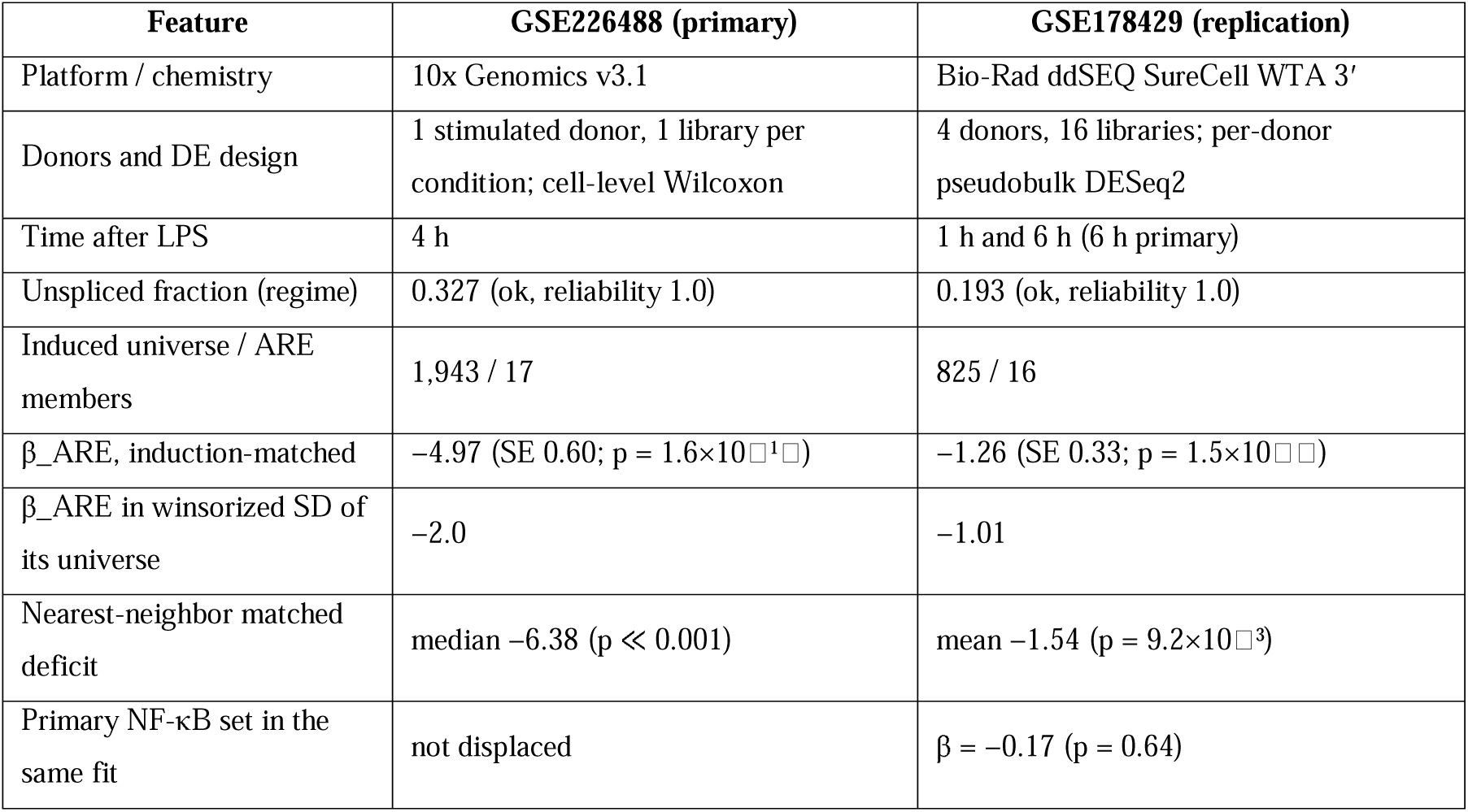
The primary and the replication LPS series. Both arms were quantified from raw FASTQ with the same STARsolo Gene+Velocyto configuration and scored with the same induction-matched procedure; β_ARE is reported both on the raw transcription_support scale and in units of the winsorized spread of its own induced universe, since the two universes differ. The coefficient rows are the differences summarized on one scale in Fig. 3H; Fig. S5A–B supply induction evidence and the pooled-design necessity.

### Soft per-gene labels and hard program-level calls implement a risk-aware annotation policy

Per-gene signal is real but modest, and headroom above it is small. The oracle upper bound—matched-abundance AUC when true labeled unspliced count is used directly as predictor—reaches only 0.68 on RPE1, against 0.61–0.63 for the residual. Per-gene AUC should be read against this low ceiling, set by sparse, noisy unspliced counts and intrinsic non-identifiability of single-gene mechanism. Thresholding behaves accordingly: at the default cut on RPE1 the transcription-driven call reached precision 0.663 and the stabilization-driven call 0.591 at 61% coverage (the 0.66 ceiling in Fig. 4A); raising the threshold (high_precision preset) halved the transcription FPR from 0.188 to 0.086 without raising precision—threshold is an FPR/recall lever, not a precision lever. Hard binary filters on the residual are therefore unsafe; on low-capture data the residual can mislabel classical immediate-early genes. scATrans keeps list membership DE-defined and exposes only soft per-gene labels, with confidence scaled by regime reliability. Candidate orthogonal validators—chromatin accessibility, ARE motif density, velocity-based reliability, Δ(u/s) decomposition—did not support genome-wide per-gene mechanism claims after appropriate controls (Fig. S3), motivating the program-level design. For zero-referenced program placement we recommend the permutation-calibrated score.

Pooling is the precision lever. Competitive Mann–Whitney tests compared transcription support to background within gene sets (BH FDR; Fig. 4B). On RPE1, separating realistically impure (70% pure) programs raised set-level AUC from 0.63 at k = 10 to 0.785 at k = 50, and to 0.960 for pure programs. Capture quality compounds the gain: on full-length K562 per-gene threshold precision reached 0.756, above the RPE1 ceiling of 0.66 (Fig. 4A), and program pooling exceeded RPE1 at every set size, reaching set-AUC 0.919 (precision 0.832, FPR 0.168) at 70% purity and k = 50. These results assume abundance- and length-matched, mechanism-coherent sets; where induction strength varies widely the induction-matched test should be primary, and per-gene mechanism_class must never feed pathway over-representation analysis (ORA) (Fig. S4B).

One consequence deserves emphasis: the residual is not a filter for removing DE false positives. Stabilization-weighted DE hits are frequently true biological changes— cytokine mRNA stabilization is the paradigm—not errors. Under modest per-gene AUC, filtering is destructive where annotation is not.

**Figure 4.**
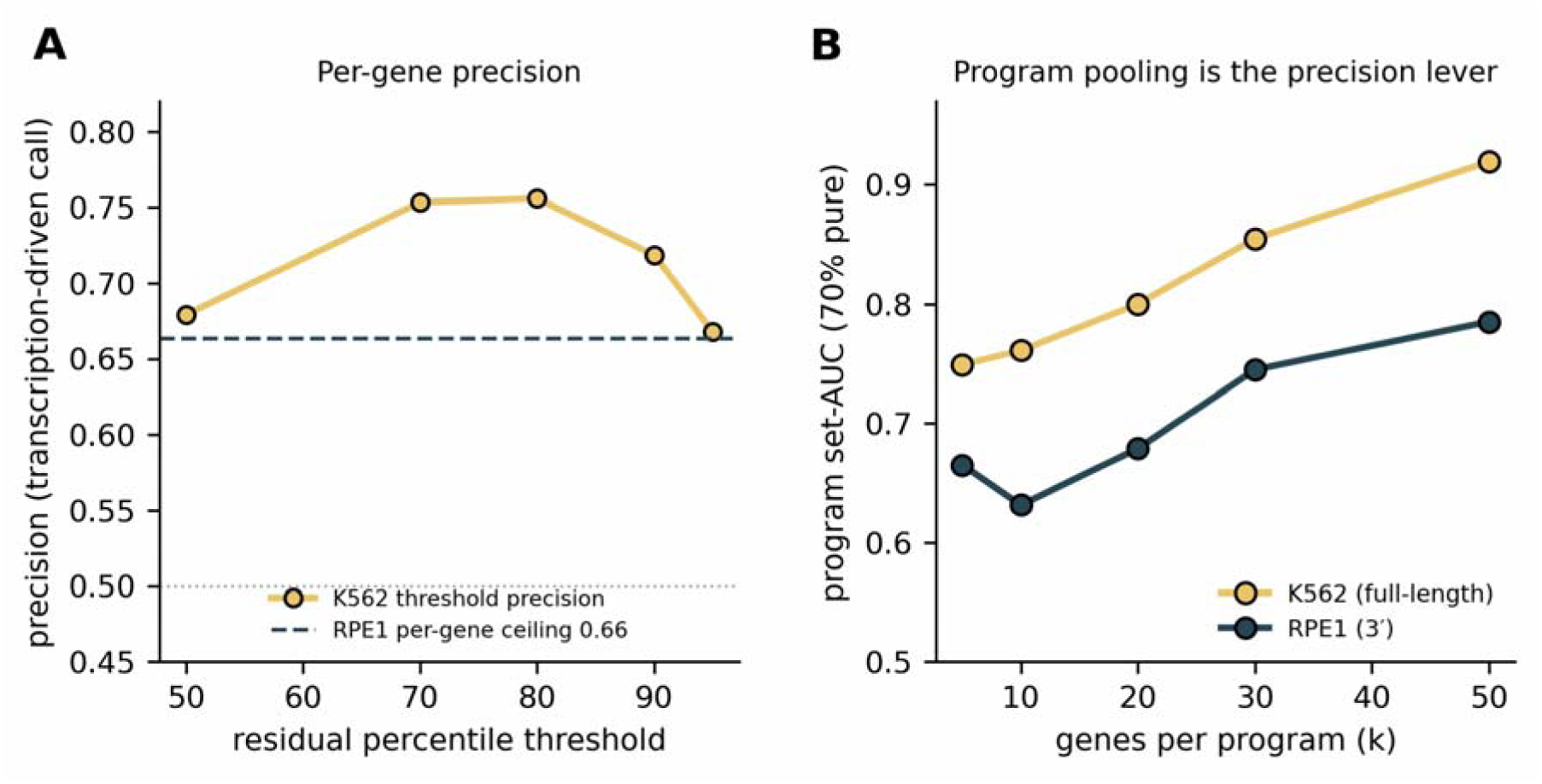
Annotation policy: per-gene ceiling versus program-level precision. (**A**) On K562, threshold precision for transcription-driven calls peaks at 0.756 (residual q70–q80), exceeding the RPE1 per-gene ceiling of 0.66. (**B**) Program pooling is the precision lever: set-level AUC rises with program size k and is consistently higher on full-length K562 than on 3′ RPE1 (synthetic 70%-pure programs). The oracle upper bound is the matched-abundance AUC when true labeled unspliced counts serve directly as predictor (≈0.68 on RPE1), bounding achievable per-gene performance.

### An optional detection score recovers active transcription on a separate axis

Length and intron residualization improve interpretability of the mechanism residual— demoting nuclear lncRNA and long-gene artifacts—but can suppress long, genuinely induced genes. scATrans therefore offers an optional nascent-detection score on a separate axis: a pseudobulk two-sample Poisson/conditional-binomial nascent z without length residualization, optionally paired with a DE-reproducibility gate. On sci-fate it raised matched-DE AUC for long active genes from 0.61 to 0.78, and the reproducibility gate eliminated DE-up false positives (0.32 → 0; Fig. 5A).

The two axes must stay decoupled. The detection score is induction-coupled; letting it drive mechanism labels collapsed the ARE stabilization call on GSE226488 (Fig. 5B)— itself evidence that the induction-normalized residual, not nascent abundance, carries the stabilization signal. Used as intended, the score refines lists rather than mechanism: restricting the GSE226488 induced universe (1,943 genes) to reproducibly nascent genes removed 557 constitutively expressed housekeeping and RNA-processing genes (ribosome biogenesis, splicing, spliceosome, proteasome; BH p < 10 ¹) while retaining the induced immune program, including 15 of 17 ARE cytokines (Fig. 5C). The removed genes are shorter, highly expressed, and lack evidence of new transcription; the effect is dataset-dependent and weaker when the response is more focused.

**Figure 5.**
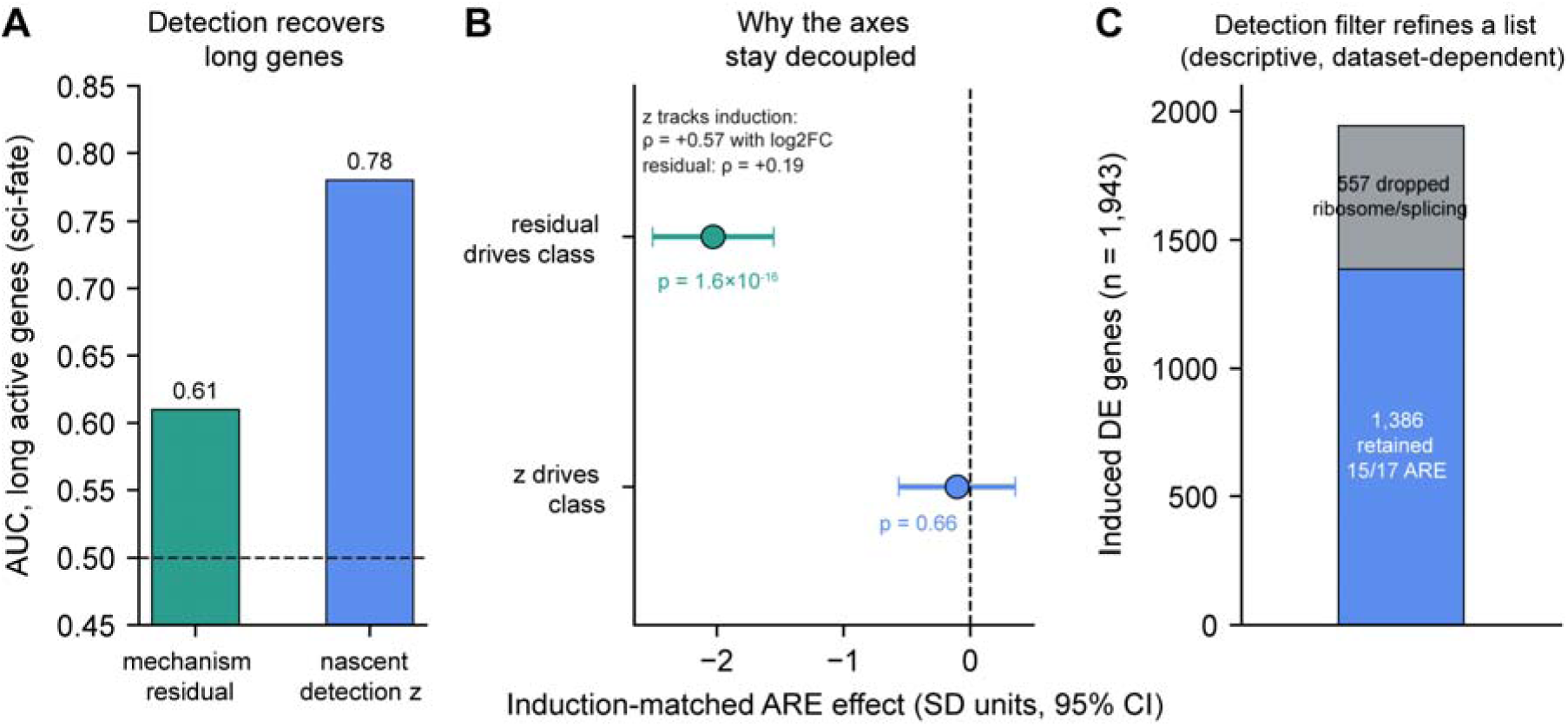
Two axes: mechanism residual versus nascent detection score. (**A**) The decoupled detection score recovers long, genuinely active genes (sci-fate matched-DE AUC 0.61 → 0.78) that length-residualized mechanism scoring suppresses. (**B**) The axes must stay decoupled. Points show induction-matched ARE effects under each axis with 95% CI, in SD units of that score’s winsorized spread over 1,943 induced genes: the stabilization call is strong when the mechanism residual defines the class (−2.0 SD; p = 1.6×10□¹) but vanishes when the induction-coupled detection z defines it (−0.10 SD; p = 0.66). Competitive Mann–Whitney agrees (p = 1.6×10□□vs. p = 0.72). The cause is coupling to induction strength: nascent z correlates with log2FC at ρ = +0.57 versus +0.19 for the residual (the +0.165 quoted in the exon–intron comparison refers to the smaller n = 1,818 universe). (**C**) Restricting 1,943 induced DE-up genes to nascent-active genes (de_reproducible and nascent z > 0) removes 557 constitutive housekeeping/RNA-processing genes while retaining the induced immune program, including 15 of 17 ARE cytokines. Descriptive refinement, not a mechanism claim.

Taken together, these results support scATrans as a mechanism annotation layer whose evidence is grounded in metabolic labeling and in known biology, whose per-gene effect size is modest and capture-dependent, and whose confident claims belong at the level of mechanism-coherent gene programs.

## Discussion

scATrans occupies a focused niche between conventional DE and RNA velocity. Relative to DE it adds a second axis—the reference-corrected unspliced residual—separating transcription-weighted from stabilization-weighted changes. Relative to velocity it favors a static condition-comparative contrast over dynamical trajectory extrapolation and strong kinetic identifiability claims ^11,12^. For mechanism annotation, this is what the data support.

The static residual recovers measured degradation γ at matched abundance as well as or better than steady-state or full dynamical velocity γ (AUC 0.627 versus 0.596 and 0.584 on RPE1; on the sparse organoid all static predictors approach chance while dynamical γ falls below it; Table 1), while remaining defined for far more genes. The bulk exon– intron comparison is convergent: on the LPS program both return the same call at the same effect size (Table S3). scATrans does not claim a superior per-gene contrast; it supplies the inference framework that turns the contrast into interpretable single-cell annotation—DE-defined membership, gene-structure residualization, capture-regime gating, induction-matched testing, and a permutation-calibrated program score whose zero is the gene set’s own expectation under label permutation. Two observations indicate these steps, not the underlying statistic, carry the weight. First, two per-gene scores matched to the residual on the γ benchmark lose the ARE call entirely at program level (Table S3)—per-gene equivalence does not imply program-level equivalence. Second, calibration behaves as an offset correction should, returning the transcriptional control toward empirical zero without moving the stabilization program off the pole (Fig. 3F). Spliced and unspliced layers are more reliably read as a static contrast than fitted as a dynamical system; the design follows: annotate mechanism at program resolution, do not claim identifiable rates.

The organizing principle is separation of tasks. DE answers which genes changed, under a statistical model chosen for the design, preferably pseudobulk. The residual asks whether the change is more consistent with elevated transcription or stabilization. The optional detection score asks which genes show elevated nascent counts, without redefining mechanism classes.

Relative to neighboring uses of the same layers, the boundary is practical. Raw unspliced abundance or a simple u/s ratio does not separate DE membership from annotation, does not residualize gene structure, and offers no induction-matched program test. RNA-velocity pipelines target trajectories and rate parameters ^9–12;^ even when they return γ, it is sparse, configuration-sensitive, and no more informative than the static residual. The bulk exon–intron contrast is the closest precedent and, as a per-gene score, an equal one; the difference lies entirely in the surrounding inference framework. Bulk frameworks decomposing synthesis and degradation from labeling time courses answer a related kinetic question but require dedicated experiments. Single-cell frameworks such as dynamo ^20^ can run on conventional splicing data, yet their distinctive output—absolute, identifiable rates—derives from reconciling splicing with metabolic-labeling kinetics, the very layers scATrans works without; on label-free data they reduce to a velocity estimator of the family already represented by the scVelo baseline (Table 1).

The biological motivation is the distinction labeling was invented to make. Transcriptional induction and mRNA stabilization are controlled by different machinery: promoter and enhancer logic on one hand; RNA-binding proteins and ARE-mediated decay on the other. Standard DE collapses both into a single fold-change. The LPS– PBMC analysis does not propose a new ARE catalog; it shows that a curated stabilization program can be recovered as stabilization-weighted, relative to NF-κB, from ordinary 10x layers after induction matching, while an independent glucocorticoid program reads transcription-weighted. The residual offers a label-free, program-level indication of which regulatory layer is more consistent with the data—useful for interpreting inflammatory and stress responses and for prioritizing follow-up, not for asserting mechanism gene by gene.

Validation clarified where the method’s value lies. The residual does not outperform DE at recovering induced genes; its contribution is mechanism partition at program resolution. The signal reproduces across two labeling technologies and three systems, scales with intron-capture quality, and is anchored in measured degradation rates. Full-length NASC-seq2 gives the strongest per-gene read (matched-abundance AUC 0.74, or 0.68 after length/intron correction; program-level set-AUC 0.92 for 70%-pure sets at k = 50), reframing the 0.66 precision ceiling on 3′ data as largely a measurement limit. That arm is not a re-run of the pulse–chase circularity concern: K562 uses orthogonal 4sU conversion chemistry and a relative labeling rate rather than a chase-derived γ from the same steady-state u/s model that defines the residual—the least circular of the three systems. Within-dataset thinning supports this: reducing K562 intronic counts 17-fold left matched-abundance AUC essentially unchanged (0.742 → 0.724), still far above 3′ RPE1 (0.627).

That inference should be stated with appropriate strength. The three-system ordering (0.54 sparse 3′ organoid, 0.63 3′ RPE1, 0.74 full-length K562, with per-gene precision rising from 0.66 to 0.76) is a cross-system comparison; thinning rules out depth but not read placement or cell type, which remain confounded in a single full-length dataset. The observation is consistent with intronic coverage across the gene body being the operative factor, but does not establish it. Protocols with better pre-mRNA recovery should improve annotation with no model change—a prediction this work motivates rather than demonstrates.

The practical consequence: scATrans recovers information otherwise obtainable mainly through metabolic labeling—whether a change is transcription- or stabilization-weighted—from counts standard scRNA-seq already produces, without 4sU or 5-EU chemistry, timed pulses, or additional libraries. It can be applied retrospectively to unlabeled public datasets wherever these layers exist, supporting program-level mechanism typing of perturbations and prioritization of candidates for confirmatory labeling. It remains a second axis on an existing DE table, better at ranking hypotheses than at replacing labeling experiments.

In practice, analysts should: (i) run regime diagnosis before interpreting any label; (ii) select genes by DE criteria appropriate to the design, preferably with biological replicates; (iii) treat per-gene mechanism classes as soft hints only; (iv) make induction-matched program tests primary unless members are already matched for induction; (v) never hard-filter DE lists by the residual; and (vi) keep detection scores and mechanism labels on separate axes (Fig. 5), never feeding soft mechanism_class into pathway ORA (Fig. S4). Because the residual is strongest early and decays as responses approach steady state (Fig. 2), acute time courses should be read at early time points and at program level.

### Limitations

**Effect size.** Per-gene effect sizes are modest (matched AUC 0.63 on scEU RPE1 against an oracle ceiling of 0.68). Claims should be framed at program level; full-length capture raises both figures.

**Regime dependence.** Low or extreme unspliced capture degrades reliability; the 0.10–0.45 trusted band is a heuristic default, not a fitted threshold. In 3′ chemistries, part of the intronic signal comes from oligo-dT internal priming on A-rich intronic regions rather than genuine nascent pre-mRNA 10,23, lowering fidelity and contributing to gene-structure bias; full-length capture is correspondingly less affected, consistent with the K562-versus-RPE1 gap.

**Direction.** The validated scope is up-regulated programs. Symmetric splitting of down-regulation into transcriptional shutdown versus destabilization is not supported at usable effect size; such genes are marked unclassified_down.

**No orthogonal per-gene validation.** Chromatin accessibility, 3′UTR ARE-motif density, velocity reliability surrogates, and Δ(u/s) decomposition all failed after appropriate controls—null after length control, or weaker than DE (Fig. S3). Length residualization can also suppress long induced genes, which the optional detection score recovers.

**Identifiability.** A stabilization-weighted label denotes induced mature abundance with lower-than-expected unspliced support—consistent with increased mRNA stability but not uniquely diagnostic, since splicing-rate differences, intron retention, processing delays, and prior cell-state history can produce the same pattern. The metabolic-labeling benchmark shares this limit: on pulse–chase arms, the static u/s ratio and measured γ are related through the same steady-state model, so those arms test recovery of model-implied γ ordering with a ceiling set by between-gene splicing-rate variation. The NASC-seq2 K562 arm partly breaks this loop via independent labeling chemistry and relative labeling rate, but no available benchmark removes all model dependence. “Zero-referenced” or “calibrated” placement means a program can be placed against what mature abundance predicts—never that a rate is estimated. At a single high-induction snapshot, per-gene labels are not fully identifiable and must not feed pathway-enrichment analyses (Fig. S4B). Label-permutation also reveals that ARE cytokines retain a structural support deficit (∼−2.9 robust-z) even under shuffled labels, reflecting their short, low-intron structure—the specific bias the permutation-calibrated score removes.

**Study design.** The primary ARE analysis rests on one LPS series with a single library per condition and single stimulated donor; p-values are exploratory. The four-donor replication (GSE178429) reproduces the call at β_ARE = −1.26 (p = 1.5×10) but at roughly half the standardized effect. Polarity and significance replicate across donors, platforms, and DE designs; magnitude is series-dependent. Time (4 h versus 6 h) and intron capture (0.327 versus 0.193) both predict attenuation. The full-length advantage rests on a single NASC-seq2 system where protocol and cell type are confounded; thinning excludes depth but leaves both unresolved.

### Outlook

Natural extensions include multi-dataset atlases of program-level mechanism calls, integration with RBP/ARE/m6A target resources for convergent checks, mechanism-of-action typing in perturbation screens, additional full-length labeling datasets to separate protocol from cell type, and cross-platform reliability estimators. Partitioning down-regulation would require systems combining measured degradation kinetics with a genuine destabilization-driven contrast; none currently available meet that bar. Because the method is capture-limited rather than model-limited, fuller-length chemistries with better pre-mRNA recovery should directly improve annotation.

## Methods

### Overview of the recommended workflow

The primary analysis route is mechanism partition of DE-selected genes:

- **Pre-flight:** compute the global unspliced fraction and map it to a reliability score and regime label (ok / low_unspliced / high_unspliced).
- **Select:** run or ingest DE; retain genes by significance and effect-size gates (defaults: BH FDR < 0.05 and |log2FC| ≥ 1, direction-aware). Prefer pseudobulk when ≥3 samples/group.
- **Annotate:** compute the reference-corrected unspliced excess, optional gene-structure residualization, robust-z transcription_support, soft mechanism_class, and confidence × reliability.

Aggregate (optional): program_mechanism over user gene sets (competitive tests on pooled transcription_support); program_mechanism_induction_matched (OLS and nearest-neighbor controls on log2FC) when induction varies; permutation-calibrated program score recommended for zero-referenced placement.

- **Detect (optional, decoupled):** nascent activity z-score; never writes mechanism_class.

Subsections follow the order quantities are defined, not computed: residual and bias correction come first as later steps reference them, but pre-flight and DE selection precede annotation in practice. Sample-level permutation is available under Mode A (Supplementary Methods S1) but not required for the primary claim. Public datasets: Supplementary Methods S7; curated gene sets: S5.

### Notation and observational units

Let T and R denote target and reference groups. For gene g, U and S denote unspliced and spliced abundances after size-factor normalization. Group means: Ū_T,g, S _T,g, Ū_R,g, S _R,g. The empirical unspliced-to-spliced ratio ρ avoids collision with kinetic parameters; under steady state ρ estimates γ/β and must not be read as a rate. Mode A (recommended) aggregates cells to pseudobulk before DE and residual means. Mode B uses cells (exploratory). Mode C uses kNN moment-smoothed cells for visualization only (Supplementary Methods S1).

Under the standard kinetic model du/dt = α − βu and ds/dt = βu − γs, steady-state u*/s* equals γ/β ^9,10^. The reference-corrected excess is an empirical deviation from fitted reference ρ, not a kinetic solution, identifying only compound γ/β rather than individual rates.

### Reference-corrected unspliced excess

The gene-wise reference ratio is shrunk toward a global background ratio ρ by additive pseudo-count shrinkage, and the excess is the vertical deviation of the target group from the reference line:

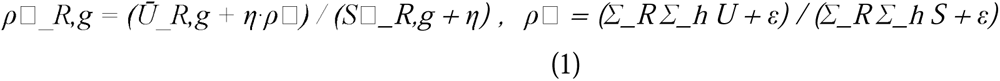

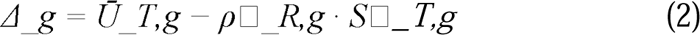

with ε = 10 and prior weight η (default 5). Positive Δ_g indicates more unspliced RNA in the target than expected from mature abundance under the reference U/S relationship. The quantity is a static, condition-comparative contrast, not a dynamical velocity estimate. Alternative ratio estimators (robust median, empirical Bayes, raw) are available (Supplementary Methods S2).

### Robust gene-structure bias correction

Gene length L_g and intron number I_g systematically influence unspliced recovery. With design vector x_g = [1, log(1+L_g), log(1+I_g)], coefficients are fitted globally by weighted Huber M-estimation ^21^ and subtracted:

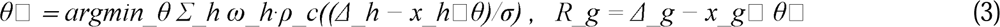

Here ρ_c is Huber loss (c = 1.35), σ a joint scale, and weights ω_h are total U+S counts winsorized at the 95th percentile. Genes lacking annotation are median-centered; failed fits fall back to global median centering and set bias_corrected = False. R_g is the bias-corrected unspliced excess used for transcription_support. Abundance-only residualization options are also implemented for sensitivity analyses. Bias correction is an empirical interpretability layer, not a claim of kinetic-rate recovery.

### DE selection (membership)

Gene-list membership is defined by DE, not by R_g. Supported backends include PyDESeq2 pseudobulk ^18,19^, Scanpy cell-level tests ^1^, mixed linear models, and optional Memento ^22^. Users may pass a precomputed DE table or a callable. Default gates (configurable): BH-adjusted p < 0.05 and |log2FC| ≥ 1, with direction split for up/down interpretation. Threshold-sensitivity grids (Jaccard versus a reference gate set) are provided for reporting. When sample_col indicates ≥3 replicates per group, pseudobulk is preferred; cell-level DE without replication is exploratory ^14–16^.

### Soft per-gene mechanism annotation

Transcription_support is a robust-z transform of the bias-corrected residual. Soft mechanism_class maps support to {transcription-driven, stabilization-driven, ambiguous, unclassified_down, unknown}, with confidence scaled by regime reliability. Labels are annotations only, never removing genes from DE lists; they denote relative transcription support among DE hits under a static residual, not absolute α /γ decomposition. A stabilization-driven label denotes induced mature abundance with lower-than-expected unspliced support—consistent with increased mRNA stability but not uniquely diagnostic, since splicing-rate differences, RNA-processing delays, and prior cell-state history can also lower unspliced support. When regime reliability is near zero, hard labels are suppressed to ambiguous. A per-gene induction-confound flag down-weights mechanism_confidence for high-induction stabilization calls (|log2FC| ≥ 90th percentile), where the static residual is non-identifiable at a single snapshot; it adds an induction_confounded column from logFC alone and leaves transcription_support and program calls unchanged.

### Program-level mechanism calls

Member-gene transcription_support is compared to background by competitive Mann– Whitney with Benjamini–Hochberg FDR^23^. Background is all other genes with defined residuals in the gene universe—the same genome-wide reference against which transcription_support is standardized. When members differ systematically in induction strength, length, or abundance, induction-matched tests are preferred. The program table is the preferred locus for binary mechanism claims. Threshold-free pooling outperforms hard gene cutoffs; separation improves with set size and intron-capture quality when sets are mechanism-coherent (see Results). For varying induction strength, induction-matched tests (support ∼ log2FC + membership; nearest-neighbor matching on log2FC) are available as program_mechanism_induction_matched and partition_de_by_mechanism(induction_matched=True).

### Permutation-calibrated program placement

Competitive and induction-matched tests are relative, ranking sets against background or matched non-members without supplying a zero—transcription_support is standardized genome-wide while each gene set has its own length, intron count, and expression composition. The calibrated program score supplies that zero empirically.

Let m be mean transcription_support of program members. Condition labels are permuted across cells, optionally within design blocks (donor, batch) to preserve condition-by-block imbalance in the null. Each replicate follows the identical chain—same DE table, residual estimator, bias correction, robust-z standardization, gene universe—yielding replicate mean m*_b. Gene-set membership is frozen (never recomputed), requiring a precomputed DE table; recomputing selection within replicates would make the null absorb DE-selection noise instead of isolating structural offset.

The calibrated displacement is: calibrated = m − mean_b(m*_b), reported with null standard deviation (quantifying offset stability, not sampling spread of m). Permutation p-value uses the (b+1)/(n+1) estimator², bounded below by 1/(n+1).

GSE226488 calibration uses n = 1,000 shuffles (four chunks of 250, pooled exactly), giving p-floor of 0.0010; primary quantities are displacement and null spread, not p-value. The sci-fate glucocorticoid calibration uses identical protocol on 2 h versus 0 h, with two design-specific differences: membership is curated rather than DE-selected (restrict_to_selected=False; 20/21 symbols pass expression filter), and cells are restricted to the two contrasted timepoints before permutation. Subsetting is inert for observed statistics (r = 1.000 versus full-series run).

The function program_mechanism_permutation_calibrated (scatrans.tl) returns n_genes, observed_mean, null_mean, null_sd, calibrated, p_perm, and n_perm_effective per program; restrict_to_selected=True intersects each program with DE-selected genes.

### Optional nascent detection score (decoupled)

When add_nascent_score=True, scATrans computes a nascent activity score from pseudobulk Poisson/conditional-binomial contrasts without length residualization; this score is for prioritization only and must not overwrite mechanism_class (see Fig. 5B).

### Regime diagnosis

The global unspliced fraction is mapped to reliability ∈ [0,1] with a U-shaped schedule: full trust in an intermediate band (default ∼0.10–0.45), declining toward extremes (low capture or nuclear/gDNA-like inflation). This score multiplies mechanism_confidence and is recorded in run metadata; the band and shape are heuristic defaults exposed as parameters.

### Software implementation

scATrans (version 0.10.9; adds program_mechanism_permutation_calibrated for calibrated placements) is pure Python on AnnData. Primary API:

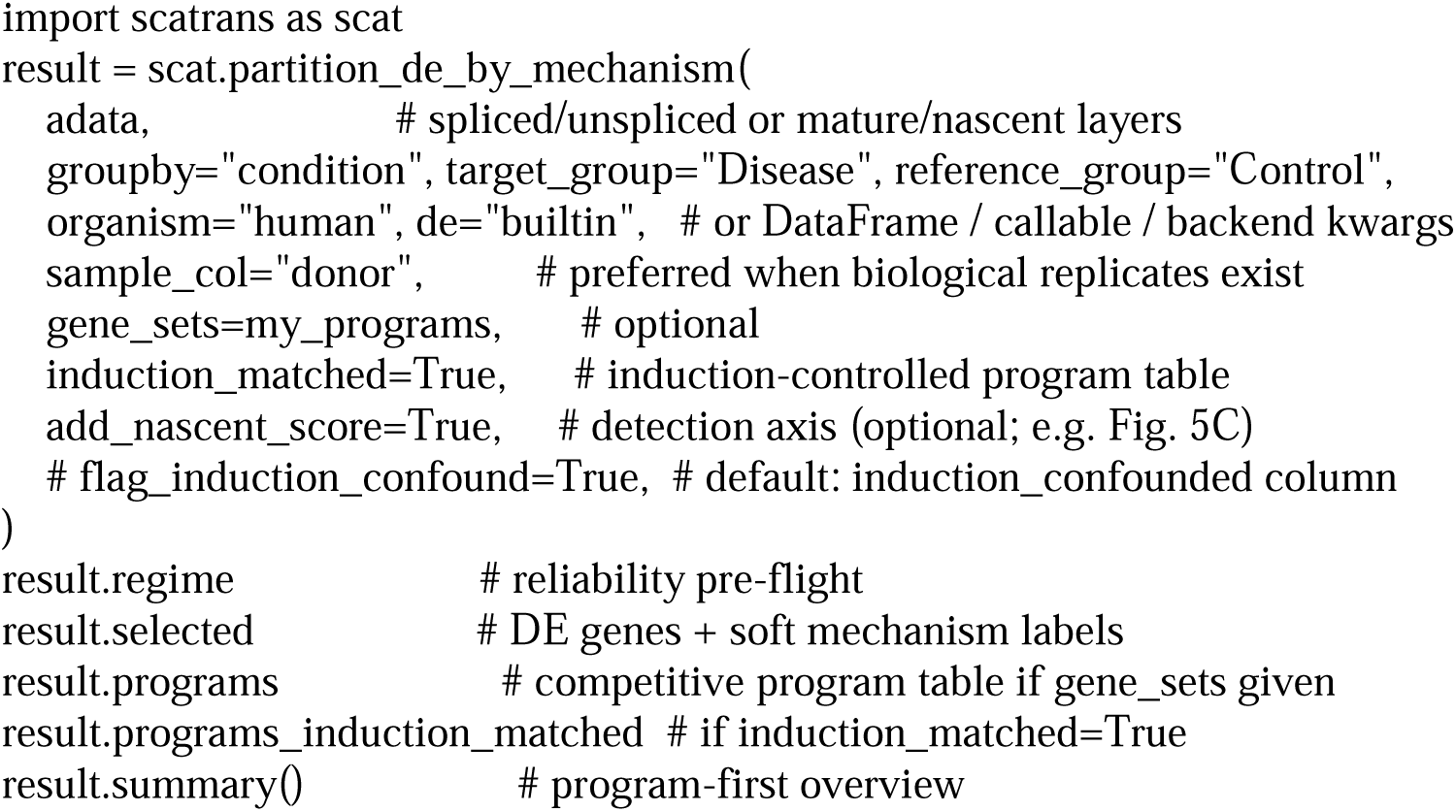

Supporting modules: regime diagnosis (scat.qc), plotting (scat.pl), enrichment (ORA/GSEA over bundled GO/KEGG; ORA on mechanism_class-stratified tables discouraged), and GTF-based gene-feature generation. Unit tests run across Python 3.9–3.12. Pre-ranked GSEA (gseapy) on transcription_support ranking was used for threshold-free program evidence (Fig. 3D). Core computation is per-gene group means plus a single global robust regression, scaling linearly in genes × cells.

### Statistics and reporting

Scale note: Three scales appear below: (i) transcription_support, a genome-wide robust-z (≈ mean 0, SD 1); (ii) induction-matched β_ARE, estimated on winsorized support within the induced universe (not genome-wide z); (iii) for cross-series comparison, β re-expressed in that universe’s winsorized SD. None should be read as Gaussian SD.

Unless noted, multiple testing uses Benjamini–Hochberg FDR ^23^, and program tests are competitive gene-set tests on transcription_support (robust-z of bias-corrected residual; genome-wide mean ≈ 0, SD ≈ 1). Three scales must be distinguished (see Results scale note): (i) robust-z is standardized genome-wide; (ii) induced-gene subsets are heavy-tailed and left-skewed (winsorized SD 2.45 on GSE226488, 1.25 on GSE178429 6 h), so OLS coefficients live on subset-specific winsorized scales; (iii) cross-series comparisons re-express coefficients in each universe’s winsorized SD. Neither scale should be read as Gaussian SD.

Induction-matched ARE analyses use: (i) OLS of 1–99% winsorized support on log2FC + ARE membership within induced genes, and (ii) nearest-neighbor matching (k = 5) on log2FC with Wilcoxon test. Winsorization is conservative (without it β_ARE = −16.8, p ≈ 10), so reported coefficients are lower bounds.

ROC-AUCs are computed within abundance deciles (γ terciles) with 1,000-replicate bootstrap 95% CIs. For Table 1, Spearman correlations use all genes passing filters (RPE1 n = 7,629; K562 n = 9,110; organoid n = 7,027), while tabulated n is the matched-abundance subset. Raw u/s ratio can lead on Spearman yet trail on matched AUC because abundance alone tracks γ (ρ = −0.46 RPE1, −0.24 K562), and matched-abundance removes that component. Organoid and RPE1 rows used identical procedures; velocity γ is undefined on K562 (single pseudobulk).

Replicate-valid robustness (Fig. S2) uses per-donor pseudobulk DESeq2 (PyDESeq2) on eight demuxlet-resolved donors, with concordance AUC scoring cell-level recovery of pseudobulk-validated induced sets. The EISA comparison normalizes intronic/exonic pseudobulk counts to CPM; two-condition contrast is Δlog2(intron) − Δlog2(exon) between LPS and resting. Both scores undergo identical winsorization, OLS, and nearest-neighbor matching as transcription_support, with coefficients divided by winsorized SD for comparability. All cutoffs, backends, and package versions are in legends and Supplementary Methods S6.

## Data and code availability

scATrans is open source under the Apache-2.0 license. This manuscript describes software version 0.10.9.

- **Package:** https://pypi.org/project/scatrans/ · Bioconda: https://anaconda.org/bioconda/scatrans
- **Source:** https://github.com/leelieber2025/scATrans
- **Documentation:** https://scatrans.readthedocs.io
- **Archived software (Zenodo):** https://doi.org/10.5281/zenodo.21365873

Public datasets are available from GEO and related repositories under the accessions listed in Supplementary Methods S7. Bundled gene-set data (GO, KEGG) carry separate licensing terms, documented in the package.

## Author contributions

Z.L. conceived the method, implemented the software, designed and performed the analyses, and wrote the manuscript. A.W.J. edited the manuscript. SX.L. implemented the software.

## Competing interests

The author declares no competing interests.

## Funding

No specific funding was received for this work. The authors used personal computational resources. We are also grateful to Sharon Onggo for reviewing the manuscript and providing valuable suggestions for grammar and readability.

## Acknowledgements

The author thanks the generators of the public datasets used for validation.

## Notes

https://github.com/leelieber2025/scATrans

https://scatrans.readthedocs.io/

### Competing Interest Statement

A.W.J. declared scientific advisory board member for Novadip LLC, consultant for Lifesprout LLC and Novadip LLC, and Editorial Board of Bone Research, Stem Cells, and The American Journal of Pathology. All the other authors declared no potential conflicts of interest.

