## Supplemental for "scATrans: annotating single-cell differential expression as transcription- or stabilization-weighted using unspliced RNA"

### **Supplementary Methods**

#### S1. Analysis routes (Modes A–C)

Mode A (pseudobulk): observational unit = biological sample; DE with PyDESeq2 by default; preferred for sample-level residual permutation when requested. Mode B (cell-level): Scanpy tests; exploratory under pseudoreplication. Mode C (kNN moment-smoothed cells): visualization of nascent excess; not for replicate-level claims. Shared core: Eqs. 1–3 of the main text.

#### S2. Optional reference-ratio estimators

Default gamma_method=heuristic_shrink (Eq. 1). Alternatives: robust_median (median anchor over expressed genes); empirical_bayes (log-ratio shrinkage with MAD prior; recommended for small reference groups; prior fixed under permutation); raw (no shrinkage).

#### S3. Optional DE backends

PyDESeq2, Scanpy t-test/Wilcoxon, mixed LM (fixed condition, random sample), Memento. When sample-level permutation of residual statistics is requested, observed and null statistics must use the same backend ^14,15,24^.

#### S4. Output field index (primary partition path)

regime / reliability; DE columns (logFC, p_adj, …); unspliced_excess_delta; unspliced_excess_residual; transcription_support; mechanism_class; mechanism_confidence; induction_confounded; optional nascent activity columns; programs table (program name, direction, p, FDR, effect); optional programs_induction_matched (when induction_matched=True); the calibrated program table from program_mechanism_permutation_calibrated (n_genes, observed_mean, null_mean, null_sd, calibrated, p_perm, n_perm_effective); meta flags such as hard-label suppression under low reliability and pseudoreplication_warning when sample_col is absent.

#### S5. Gene sets and ARE curation (GSE226488)

Design. GSE226488 is a CNRGH 10x v3.1 PBMC series (resting, LPS 4 h, fresh and frozen libraries). This manuscript uses only GSM7077865 (resting, replicate 1) and GSM7077866 (LPS 4 h, replicate 1). FASTQs were reprocessed with STARsolo Gene+Velocyto on GRCh38; cells were restricted to Gene/filtered barcodes; AnnData layers were spliced, unspliced, and ambiguous with X = spliced (15,449 cells × 62,754 genes). Regime diagnosis reported unspliced fraction 0.327 (ok, reliability 1.0). DE used the built-in cell-level Wilcoxon backend without sample_col (no pseudobulk), because only one library exists per condition; DE p-values and program competitive p-values are therefore exploratory under pseudoreplication ^14–16^. No cell-type annotation or monocyte subsetting was applied; results are whole-PBMC mixture effects.

transcription_support scale. In this run, transcription_support is a robust-z transform of the length/intron bias-corrected unspliced excess residual (Pearson correlation with the residual = 1; genome-wide SD ≈ 1). That unit spread is genome-wide; the induced-gene universe used for matched tests is a heavy-tailed, strongly left-skewed subset of it (winsorized SD 2.45), which is why coefficients estimated on that universe are reported in units of its own spread and why robust-z units should never be read as normal standard deviations. Extreme negative outliers among highly induced cytokines (e.g. CCL4) are genuine on the residual scale; induction-matched OLS therefore winsorizes support at the 1st and 99th percentiles of the induced-gene universe before fitting support ~ log2FC + is_ARE.

ARE / ZFP36 set (expected stabilization-weighted). Literature classic TTP/ZFP36 targets and AU-rich inflammatory cytokines/chemokines ^2^, plus closely related LPS-induced ARE-bearing mRNAs used as a coherent post-transcriptional program: TNF, IL6, CXCL8, IL8 (alias of CXCL8), PTGS2, IL1B, IL1A, CXCL1, CXCL2, CXCL3, CCL2, CCL3, CCL4, CCL3L1, CSF2, CSF3, IL10, ZFP36, PTX3, SOD2, PLAUR. This is 21 symbols but 20 distinct genes, since IL8 is retained only as an alias check for CXCL8; counts reported elsewhere as “21” refer to symbols.

NF-κB primary transcriptional set (relative control). IκB-family feedback regulators, NF-κB family members, and selected immediate-early transcriptional components regulated primarily by transcription rather than by ARE decay: NFKBIA, NFKBIZ, NFKBIE, NFKB1, NFKB2, RELB, REL, TNFAIP3, TNIP1, TNIP2, BIRC3, NFKBIB, EGR1, EGR2, JUNB, ATF3, IER2, IER3, GADD45B (19 symbols).

Exclusions and dual regulation. The two lists are constructed to be disjoint (empty intersection). Genes known to be both strongly transcriptionally induced and ARE-bearing were not dual-listed; when a symbol appeared in both candidate pools during curation it was assigned to at most one set. ZFP36 (TTP) itself is down-regulated in this LPS-versus-resting table and is therefore excluded from induced-universe matched tests, though it remains in the deposited ARE list. IL8 is retained only as an alias check for CXCL8; CCL3L1 may be absent from the STARsolo gene index. An optional positive-control TF set used in secondary scripts (IRF/STAT axis): NFKB1, NFKB2, RELB, REL, IRF1, IRF7, STAT1, STAT2, IRF9, NFKBIA, SP100, BATF, BCL3.

Primary numerical results (locked). Selected DE genes (padj < 0.05, log2FC ≥ 1): 803. Induced universe for matching: 1,943. Program call (partition_de_by_mechanism): ARE n = 16, mean support −17.45 (median −8.46) vs background +0.54 (median +0.29), p = 1.58×10⁻⁸; NF-κB primary n = 10 in the selected table, mean support −0.38 (median −0.76; with n = 10 this central value is imprecise and is reported for completeness rather than as a test). Background here is the DE-selected set minus both curated lists (n = 777); interquartile ranges are [−32.25, −1.70] for ARE, [−1.74, −0.18] for the NF-κB set and [−0.44, +1.40] for background, and the ARE-minus-background difference computed on unrounded medians is −8.74 (Hodges–Lehmann −9.92). Matched OLS: β_ARE = −4.97 (SE 0.60), p = 1.62×10⁻¹⁶; β_logFC = +0.038 (p = 0.21). Nearest-neighbor median (ARE − matched non-ARE support) = −6.38. Gene-set text files: paper_analysis/05_gse226488_ARE/deposited/. Of the 21 curated ARE symbols, 19 were present in the STARsolo gene index for this library, 17 had finite transcription_support in the induced universe used for matching, and 16 fell within the DE-selected program (padj < 0.05, log2FC ≥ 1); the differing counts across analyses reflect these successive filters, not different gene sets. Robustness of the matched OLS (deposited/ARE_robustness_loo_curation.txt): leave-one-ARE-out β_ARE ranges [−5.32, −4.75] (all p < 2×10⁻¹⁴; largest shift 0.36 when PLAUR is dropped; removing the extreme-support outlier CCL4 gives β = −4.76). Alternate ARE curations give β_ARE = −4.89 (drop SOD2/PTX3/PLAUR, n = 14), −4.90 (drop those + PTGS2, n = 13), −5.19 (cytokine/chemokine-only, n = 12), and −5.74 (canonical TTP core: TNF, IL6, CXCL8, IL1B, CXCL1, CXCL2, CCL3, CSF2; n = 7). The induction-matched stabilization call is therefore stable across single-gene removal and curation choice. Scale reference for the coefficient (induced universe, n = 1,943): transcription_support percentiles p1 = −7.57, p5 = −2.73, p25 = −0.71, median = −0.01, p75 = +0.86, p95 = +4.23, p99 = +11.55 (range −75.2 to +26.3); winsorized SD 2.45; IQR 1.56; β_ARE = −4.97 is 2.0 winsorized SD, 3.2 IQR, 26% of the 1–99% span, and moves a gene from the median to the 2.2nd percentile. EISA comparison (Table S3; paper_analysis/05_gse226488_ARE/EISA_induction_matched.py, outputs in deposited/): common universe n = 1,818, i.e. 1,943 induced genes minus 125 without a defined exon–intron contrast (exon ≥ 5 and intron ≥ 1 in the case sample), which removes one ARE member, CCL2, leaving 16; on that universe transcription_support gives β_ARE = −4.73 (SE 0.64, p = 2.0×10⁻¹³) against −4.97 on the full induced universe. Permutation calibration (paper_analysis/05_gse226488_ARE/calibration_aligned.py + combine_calibration_chunks.py; 1,000 shuffles as four seeded chunks of 250) is run on this same locked selection, so that every calibrated value refers to the 803-gene universe above: ARE n = 16, observed −17.454, null −2.938 (SD 0.128), calibrated −14.516, p_perm = 0.0010; NF-κB primary n = 10, observed −0.382, null −0.148 (SD 0.028), calibrated −0.234, p_perm = 0.0010.

#### S6. Reproducibility checklist for Results figures

- scatrans.__version__ and extras (pseudobulk, gsea, …)
- Genome build, GTF, whitelist, STARsolo/STAR parameters
- DE backend, sample_col, padj and logFC gates
- Bias method; whether add_nascent_score
- Random seeds for any permutation/subsampling
- Script or notebook path under paper_analysis/
- Plotted values with per-value provenance (paper_analysis/07_figures_and_manuscript/figure_data/figure_values.csv)

#### S7. Datasets used for validation (public)

No new sequencing was generated for this study. Validation used:

scEU-seq RPE1 pulse–chase (GSE128365): author-processed labeled/unlabeled × spliced/unspliced matrices; per-gene kinetics and matched-abundance specificity (regime unspliced fraction 0.16) ^7^.

scEU-seq mouse intestinal organoid (GSE128365): author-processed matrices; directional replication of the specificity result 8, and the second system of the Table 1 kinetic-model comparison (per-cell spliced/unspliced layers plus pulse–chase γ, so velocity estimators are defined).

NASC-seq2 K562 (full-length 4sU): raw FASTQ reprocessed with STAR (72.0% uniquely mapped) and velocyto (unspliced fraction 0.28); relative degradation axis γ_rel = −ln(1−f_new) from the new/old molecule tables; matched-abundance specificity, threshold precision, and program pooling ^6^.

scNT-seq KCl neurons (GSE141851): FASTQ reprocessing with STARsolo where applicable; new/old layers as labeling truth ^4^.

sci-fate A549 dexamethasone (GSE131351): combinatorial-indexing time course; labeling truth for the kinetic decay analysis, and a program-level mechanism case (the glucocorticoid-receptor program reads transcription-weighted at 2 h, the opposite polarity to the GSE226488 ARE program; permutation-calibrated displacement +0.092 against a null expectation of +0.430, 1,000 shuffles) ^5^.

GSE226488 PBMC resting vs LPS 4 h (10x v3.1): STARsolo Gene+Velocyto (GRCh38); Gene/filtered barcodes; cell-level Wilcoxon DE (exploratory; one library per condition); regime pre-flight and ARE mechanism biology (Tables S1–S2; S5) ^25^.

Kang IFN-β PBMC (GSE96583, 10x 3′ v1): re-aligned to GRCh38 (bamtofastq → STARsolo Gene+Velocyto); regime pre-flight ok (unspliced fraction 0.165); eight demuxlet-resolved donors enabling per-donor pseudobulk DESeq2 (eight biological replicates per condition) as a replicate-valid robustness analysis of the cell-level DE workflow (Fig. S2) ^17^.

GSE178429 LPS PBMC, four donors (Bio-Rad ddSEQ SureCell WTA 3′; the scRNA-seq leg of the Kartha et al. multi-omics superseries GSE178431, whose ATAC leg supplied the chromatin negative control): raw FASTQ reprocessed with STARsolo (soloType CB_UMI_Complex; three-block barcode resolved against the TATGCATGAC and CAGTCAGTCA linkers with an 8 nt leading UMI) plus Velocyto features on GRCh38, matching the GSE226488 configuration; control versus LPS at 1 h and 6 h × Donor 1–4 = 16 libraries, 13,438 cells, regime unspliced fraction 0.193; per-donor pseudobulk DESeq2 with donor as the sample unit, supplying the independent multi-donor replication of the ARE stabilization call.

Orthogonal/negative explorations: the scATAC/multiome leg of the same Kartha LPS superseries supplied the chromatin negative control ^26^.

Accessions, GSM IDs, gene-set files, and figure scripts for GSE226488 are summarized in Tables S1–S3 and Supplementary Methods S5 (also paper_analysis/05_gse226488_ARE/). Genome builds, whitelists, and command lines for all datasets are provided under paper_analysis/ in the source repository. Tutorial subsets (downsampled h5ad) are documented in the package tutorials.

### Supplementary Tables

**Table S1.** GSE226488 analysis design for the LPS–PBMC mechanism case study.

| **Item** | **Value** |
| --- | --- |
| GEO series | GSE226488 (CNRGH PBMC resting / LPS / fresh / frozen series) |
| Libraries used | GSM7077865 resting (rep. 1); GSM7077866 LPS 4 h (rep. 1) |
| Chemistry / platform | 10x Genomics 3′ v3.1; Illumina NovaSeq 6000 |
| Reprocessing | STARsolo Gene+Velocyto; GRCh38; called cells = Gene/filtered barcodes |
| Cells × genes | 15,449 × 62,754 (resting 7,170; LPS 8,279) |
| Unspliced fraction / regime | 0.327 / ok (reliability 1.0) |
| DE backend (this case) | Cell-level Wilcoxon (Scanpy); no sample_col / not pseudobulk |
| DE gates for selected set | padj < 0.05 and log2FC ≥ 1 (LPS vs resting) → 803 genes |
| Induced universe (matched tests) | log2FC > 0 and padj < 0.05 with finite support → 1,943 genes |
| transcription_support scale | Robust-z of bias-corrected unspliced residual (genome-wide SD ≈ 1) |
| Cell types | Whole PBMC mixture; no annotation-based subsetting |
| Biological replicates | One library per condition → exploratory DE / program p-values |

**Table S2.** Curated gene sets for GSE226488 (disjoint lists; see S5 for sources and exclusions).

| **Set** | **n listed** | **Role in test** | **Member symbols** |
| --- | --- | --- | --- |
| ARE / ZFP36 stabilized | 21 | Expected lower transcription_support (stabilization-weighted) | TNF, IL6, CXCL8/IL8, PTGS2, IL1B, IL1A, CXCL1–3, CCL2–4, CCL3L1, CSF2, CSF3, IL10, ZFP36, PTX3, SOD2, PLAUR |
| NF-κB primary transcription | 19 | Primary transcriptional / feedback controls (higher support than ARE) | NFKBIA, NFKBIZ, NFKBIE, NFKB1, NFKB2, RELB, REL, TNFAIP3, TNIP1, TNIP2, BIRC3, NFKBIB, EGR1, EGR2, JUNB, ATF3, IER2, IER3, GADD45B |

**Table S3.** EISA versus the scATrans residual under the induction-matched program test (GSE226488 LPS–PBMC). Common induced universe: n = 1,818 genes with both scores defined; 16 ARE/ZFP36 members. Scores winsorized at 1st/99th percentiles; β_ARE is the ARE-membership OLS coefficient (log2FC + ARE membership), reported raw and in winsorized SD units for cross-score comparability. NN = median difference between each ARE gene and five log2FC-matched non-ARE genes (one-sided Wilcoxon). Naive column omits log2FC covariate to isolate what induction matching changes.

| **Score** | **naive β_ARE (SD)** | **matched β_ARE (SD)** | **p (matched)** | **NN median (SD)** | **p (NN)** |
| --- | --- | --- | --- | --- | --- |
| **scATrans transcription_support** | −4.61 (−1.83) | −4.73 (−1.87) | 2×10⁻¹³ | −4.99 (−1.98) | 7.6×10⁻⁵ |
| EISA two-condition contrast (ΔI − ΔE) | −1.79 (−2.32) | −1.42 (−1.85) | 7.9×10⁻¹⁴ | −1.19 (−1.55) | 3.8×10⁻⁴ |
| EISA transcription leg only (ΔI) | +1.73 (+1.38) | −0.13 (−0.10) | 0.60 | +0.16 (+0.13) | 0.68 |
| EISA intronic level, case sample (Table 1) | −0.06 (−0.02) | −0.22 (−0.09) | 0.74 | −1.37 (−0.53) | 0.35 |

**Table S4.** Monocyte-subset and permutation validation of the ARE stabilization call (GSE226488 LPS–PBMC). Monocytes identified by lineage markers (CD14, LYZ, FCN1, S100A8/9) via Leiden clustering on spliced expression; ARE partition re-run within 3,893 monocytes (25% of PBMC; LPS 2,047, resting 1,846). The stabilization call strengthens on every test and remains far stronger than NF-κB control, ruling out cell-composition artifacts. Gating: 4 of 18 clusters with monocyte-marker score > 0.5. A looser lineage-argmax rule (5,422 cells, 35%) yields the same call (β_ARE = −7.25), confirming insensitivity to gating threshold.

| **Test** | **Whole PBMC** | **Monocytes only** |
| --- | --- | --- |
| ARE competitive mean support | −17.45 (p = 1.6×10⁻⁸) | −24.35 (p = 5.9×10⁻¹⁰) |
| Induction-matched β_ARE | −4.97 (p = 1.6×10⁻¹⁶) | −8.13 (p = 7.7×10⁻³²) |
| Nearest-neighbor median Δ | −6.38 (p ≪ 0.001) | −8.78 (p = 2.9×10⁻⁴) |
| NF-κB control (induction-matched) | not significant | β = −1.89, p = 0.014 (≪ ARE) |

### Supplementary Figures

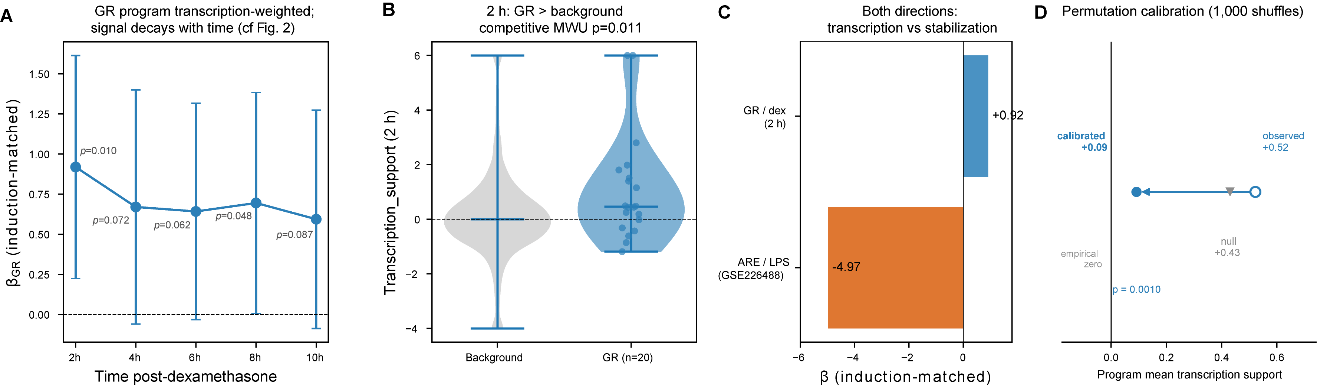

**Figure S1. sci-fate A549 + dexamethasone:** GR transcriptional program is transcription-weighted (supplementary to GSE226488 ARE stabilization case). (**A**) Induction-matched β_GR (OLS transcription_support ~ log2FC + GR-membership; 95% CI) across timepoints: positive throughout, strongest at 2 h (β = +0.92, p = 0.010), decaying thereafter consistent with residual's nascent time decay (Fig. 2). (**B**) At 2 h, transcription_support for 20 GR genes versus background (competitive Mann–Whitney p = 0.011, one-sided; direction pre-specified, unlike two-sided ARE test of Fig. 3B). (**C**) Opposite polarity: GR/dex transcription-weighted (β = +0.92) versus ARE/LPS stabilization-weighted (β = −4.97). (**D**) Permutation calibration as in Fig. 3F: observed mean (open circle), expectation over 1,000 shuffles (gray triangle), calibrated displacement (filled circle). Most raw positive offset is structural (shuffled expectation +0.43 vs. observed +0.52), yet calibrated displacement stays positive at +0.09 (p = 0.0010, floor at this replicate count)—the transcription-side mirror of ARE calibration in Fig. 3F. GR genes independently confirmed transcriptionally induced by sci-fate labeling (new-RNA log2FC ≈ +1.5–1.8); residual computed without the label.

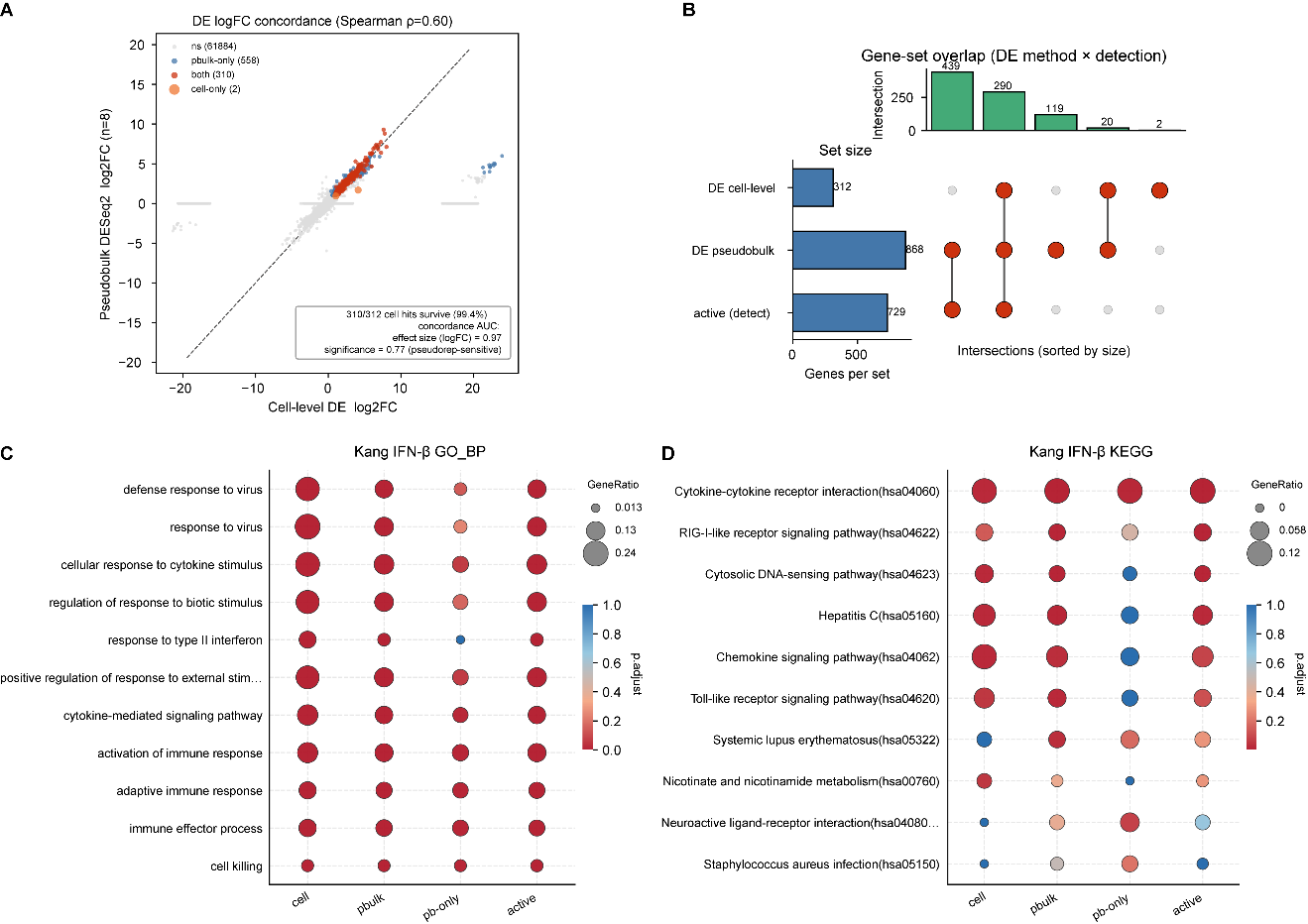

**Figure S2. Mechanism and detection calls are robust to DE replicate design (Kang IFN-β PBMCs, GSE96583; eight demuxlet-resolved donors).** (**A**) Per-gene log2FC: cell-level DE (x) versus per-donor pseudobulk DESeq2 (y); Spearman ρ = 0.60. Of 312 cell-level–selected genes, 310 (99.4%) survive replicate-valid design; concordance AUC 0.97 by effect size, 0.77 by signed significance (pseudoreplication-sensitive). (**B**) UpSet plot: cell-level DE, pseudobulk DE, and detection axis (de_reproducible and nascent z > 0). (**C, D**) GO-BP and KEGG over-representation across both DE methods, pseudobulk-unique genes (pb-only), and active set; dot size = gene ratio, color = adjusted p. Both DE methods enrich identical interferon/antiviral program; pb-only genes carry weaker secondary immune signal; active axis retains antiviral core.
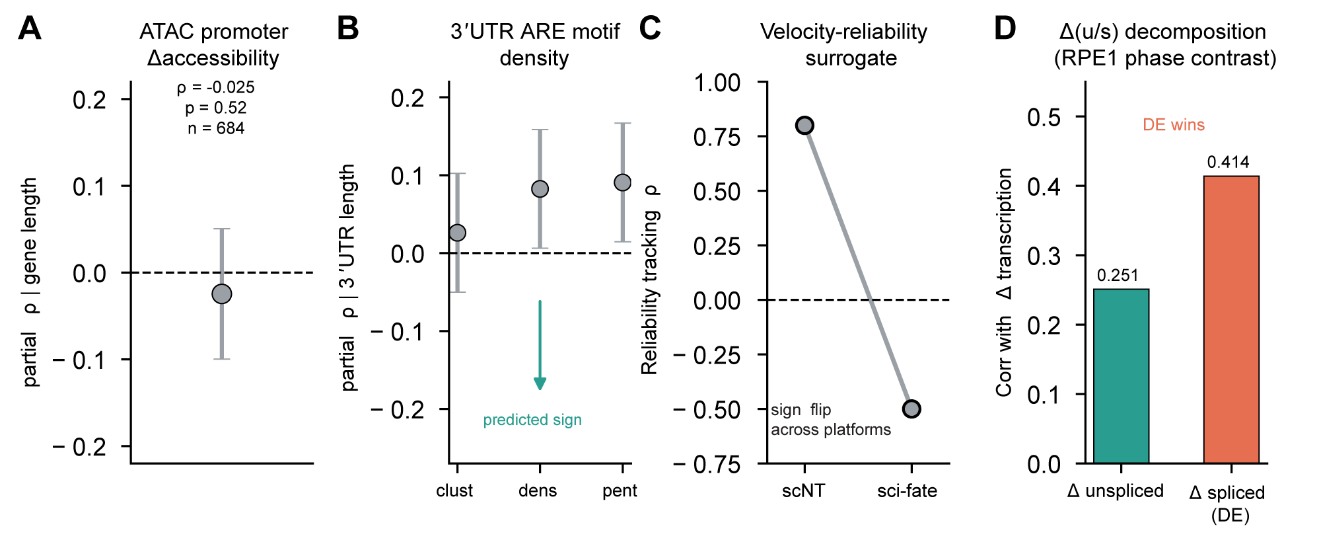

**Figure S3. Failed alternatives delimiting the claim.** (**A**) Chromatin: partial Spearman between transcription_support and LPS-induced promoter accessibility change, controlling for log gene length, ρ = −0.025 (p = 0.52, n = 684); accessibility metric valid (positive control p = 1.5×10⁻⁵). (**B**) Sequence: partial Spearman between transcription_support and three 3′UTR ARE measures (clusters, density/kb, pentamer count), controlling for log 3′UTR length (ρ = +0.03 to +0.09, n = 665). Stabilization-weighted genes should carry more ARE (negative correlation expected); all estimates are small and opposite-signed. (**C**) Label-free reliability surrogate: ρ = +0.80 on scNT-seq but −0.50 on sci-fate—does not transfer across platforms. (**D**) Δ(u/s) decomposition on RPE1 cell-cycle contrast: unspliced change tracks labeled transcription worse than spliced change (r = 0.251 vs. 0.414), so DE wins the task decomposition was designed for. These delimiters support reading the residual as static annotation rather than orthogonally validated per-gene classifier.

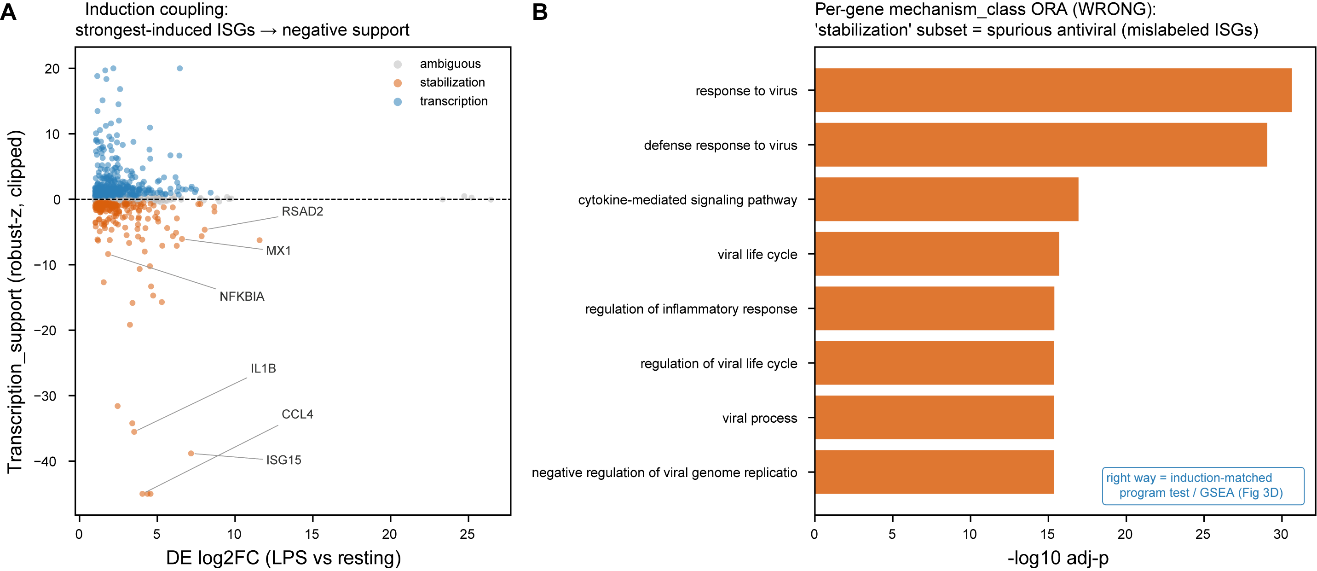

**Figure S4. The induction-confound trap: per-gene mechanism_class ORA is not pathway biology (GSE226488 LPS–PBMC).** (**A**) transcription_support versus log2FC for 803 DE-selected genes, colored by soft mechanism_class. Strongly induced ISGs (ISG15, MX1, RSAD2), NF-κB feedback gene NFKBIA, and ARE cytokines (IL1B, CCL4) all fall to negative support at 4 h—illustrating induction coupling at the gene level. The NF-κB program still sits above ARE (Fig. 3B); absolute placement requires induction matching (Fig. 3C) or permutation calibration (Fig. 3F). (**B**) ORA on the 'stabilization' subset spuriously recovers antiviral GO terms (response to virus; p < 10⁻²⁹), driven by mislabeled ISGs rather than ARE biology. Correct analyses: induction-matched program test (Fig. 3C) and threshold-free GSEA on transcription_support ranking (Fig. 3D). Enrichment: hypergeometric ORA, GO-BP, expressed-gene universe, BH FDR; p-values exploratory (one 10x library per condition; Table S1).

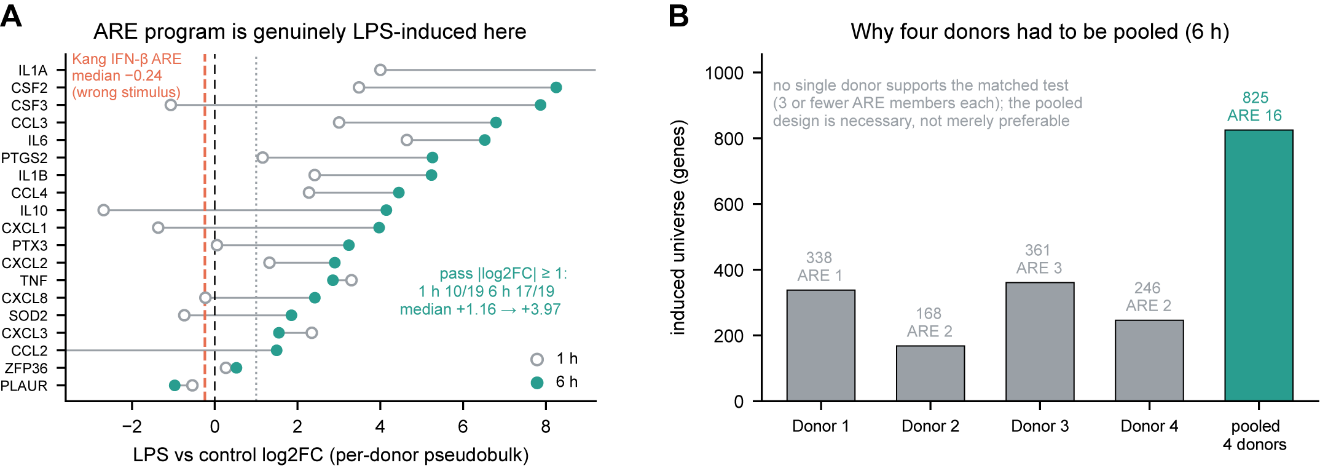

**Figure S5. Four-donor ARE replication details (GSE178429 LPS PBMC; main-text Fig. 3G, H).** Control vs. LPS at 1 h and 6 h, four donors (16 libraries, 13,438 cells; unspliced fraction 0.193), STARsolo Gene+Velocyto, per-donor pseudobulk DESeq2. (A) ARE members are genuinely LPS-induced: log2FC of 19 curated members, 1 h (open) vs. 6 h (filled); 10/19 pass |log2FC| ≥ 1 at 1 h, 17/19 at 6 h (median +1.16 → +3.97). Dashed red line: Kang IFN-β ARE median (−0.24), where this set is not induced—wrong stimulus. (B) Why donors were pooled: each donor alone yields ≤3 ARE members at 6 h, insufficient for induction-matched test; pooling gives 825 genes and 16 ARE members. Matched regression and cross-series β: Fig. 3G, H.
